# Tracking the PROTAC degradation pathway in living cells highlights the importance of ternary complex measurement for PROTAC optimization

**DOI:** 10.1101/2023.01.11.523589

**Authors:** Martin P. Schwalm, Andreas Krämer, Anja Dölle, Janik Weckesser, Xufen Yu, Jian Jin, Krishna Saxena, Stefan Knapp

**Affiliations:** Institut für Pharmazeutische Chemie, Goethe-University Frankfurt, Biozentrum, Max-von-Laue-Str. 9, 60438 Frankfurt am Main, Germany; Structural Genomics Consortium, Goethe-University Frankfurt, Buchmann Institute for Life Sciences, Max-von-Laue-Str. 15, 60438 Frankfurt am Main, Germany; Mount Sinai Center for Therapeutics Discovery, Departments of Pharmacological Sciences, Oncological Sciences and Neuroscience, Tisch Cancer Institute, Icahn School of Medicine at Mount Sinai, New York, NY 10029, USA; German Cancer Consortium (DKTK)/German Cancer Research Center (DKFZ), DKTK site Frankfurt-Mainz, 69120 Heidelberg, Germany

**Keywords:** PROTAC, Degrader, Kinetics, WDR5, live cells

## Abstract

The multistep PROTAC (PROteolysis TArgeting Chimeras) degradation process poses challenges for their rational development, as rate limiting steps determining PROTAC efficiency remain largely unknown. Moreover, the slow throughput of currently used endpoint assays does not allow the comprehensive analysis of larger series of PROTACs. Here we developed cell-based assays using NanoLuciferase and HaloTags, that allow measuring PROTAC induced degradation and ternary complex formation kinetics and stability in cells. Using PROTACs developed for degradation of WDR5, the characterization of the mode of action of these PROTACs in the early degradation cascade revealed a key role of ternary complex formation and stability. Comparing a series of ternary complex crystal structures highlighted the importance of an efficient E3-target interface for ternary complex stability. The developed assays outline a strategy for the rational optimization of PROTACs using a series of live cell assays monitoring key steps of the early PROTAC induced degradation pathway.

**Significance:** The multistep PROTAC induced degradation process of a POI poses a significant challenge for the rational design of these bifunctional small molecules as critical steps that limit PROTAC efficacy cannot be easily assayed at required throughput. In addition, the cellular location of the POI may pose additional challenges as some cellular compartments, such as the nucleus, may not be easily reached by PROTAC molecules and the targeted E3 ligases may not be present in this cellular compartment. We propose therefore a comprehensive assay panel for PROTACs evaluation in cellular environments using a sensor system that allows continuous monitoring of the protein levels of the endogenous POI. We developed a cell line expressing WDR5 from its endogenous locus in fusion with a small sequence tag (HiBIT) that can be reconstituted to functional NanoLuciferase (NLuc). This system allowed continuous monitoring of endogenous WDR5 levels in cells and together with HaloTag system also the continuous monitoring of ternary complex (E3, WDR5 and PROTAC) formation. As this assay can be run at high throughput, we used this versatile system monitoring three diverse chemical series of WDR5 PROTACs that markedly differ in their degradation properties. Monitoring cell penetration, binary complex formation (PROTAC-WDR5 and PROTAC-VHL) as well as ternary complex formation we found that PROTAC efficiency highly correlated with synergy of ternary complex formation in cells. This study represents a first data set on diverse PROTACs studying this property *in cellulo* and it outlines a strategy for the rational optimization of PROTACs. It also provided kinetic data on ternary complex assembly and dissociation that may serve as a benchmark for future studies utilizing also kinetic properties for PROTAC development. Comparative structural studies revealed larger PROTAC mediated interaction surfaces for PROTACs that efficiently formed ternary complexes highlighting the utility of structure based optimization of PROTAC induced ternary complexes in the development process.

## Introduction

Recently, PROteolysis TArgeting Chimeras (PROTACs) have gained considerable attention as new modalities for drug development as well as tools in cell biology and target validation studies.^1-3^ PROTACs rely on the ubiquitin-proteasomal degradation system by recruiting designated target proteins of interest (POI) to E3 ligases using chimeric small organic molecules that on one hand bind to the E3 ligase (handle) and on the other hand to the POI (warhead). For the development of PROTACs, the linker as connection between the E3 ligand and the warhead binding to the POI is crucial for the efficient binding of PROTACs and the orientation of both proteins in PROTAC induced ternary complexes. Identification of most suitable attachment points and linkers sometimes requires extensive optimization by medicinal chemistry methods as well as efficient screening strategies to develop highly potent degraders.^4,5^ However, as the ubiquitin degradation pathway is a complex multistep process, even for well-designed chimeric PROTACs that efficiently penetrate cells and tightly interact with both binding partners in a stable ternary complex, the effective degradation of the POI is not guaranteed as the degradation process requires also the effective transfer of ubiquitin to an available lysine at the POI as well as recruitment and effective degradation by the proteasome.^6^ To date, the mechanistic reasons leading to inefficient degradation despite tight interaction of POI and E3 ligands are usually not known and there are very few studies that also consider kinetic aspects of the PROTAC induced degradation cascade. Reviewing the literature on this topic, we realized that most studies only monitor the endpoint of degradation, thus simply analyzing the degradation of the POI by Western blotting (WB), a method with a very limited dynamic range and throughput. Additionally, most published studies have been carried out using lysed cells or biochemical interaction assays. These studies therefore do not consider the cellular environment and architecture such as the localization of the endogenous POI and its integration into large protein complexes that may restrict access to E3 ligases.

To study protein levels of a specific POI in live cells, fusion with a luminescent protein can be used allowing to trace changes in protein levels by fluorescence resonance transfer.^7^ However, attachment of tags with a considerable size may change cellular localization, stability and expression levels of the POI. To overcome this limitation, the development of the NanoBIT system has established a sensor system by an in frame fusion with a small sequence tag of only 11 amino acids, called HiBIT. This peptide has high affinity towards the LgBIT which marks the residual scaffold of the NanoLuciferase (NLuc) and complementation allows bioluminescence detection. This minimally invasive tagging for monitoring of live cell protein levels promises a tagged POI with native like behaviour.^8^ Since PROTACs must efficiently act on the initial steps in the ubiquitin degradation process, assaying each step can help to identify limiting factors of PROTAC induced POI degradation. Finally, probing the PROTAC-induced degradation of the POI yields crucial parameters for PROTAC characterization and evaluation including DC_50_ and D_max._ that cannot be accurately determined by biochemical methods. The DC_50_ describes the effective concentration required to reach 50% of the PROTACs degradation efficiency, while the D_max_ indicates the maximum percentage of targeted protein degradation. Since both values are in direct relation to each other, measurement of degradation kinetics can reveal the interdependence of D_max_ and DC_50_ together with their time dependencies allowing a deeper understanding of the degradation mechanism of action (MOA).^9^

In this study we chose chemically diverse series of PROTACs targeting the chromatin-associated WD40 repeat domain protein 5 (WDR5) as a model system to understand critical steps in POI degradation (Figure 1 A). WDR5 represents a well-studied adapter protein for the induction of protein-protein interactions (PPI). WDR5 consists of a 7-bladed β-propeller structure^10^ and is a protein which nearly exclusively consists of a WD40 domain with an N-terminal extension and therefore does not have an active site or enzymatic activity making it an interesting target for targeted protein degradation.^11^ Its main role is based on a scaffolding function in large protein complexes including NSL complex (acetyl-transferase)^12^, NuRD (chromatin remodeling and deacetylation)^13^ or the SET1/MLL complex (methyltransferase).^14^ Due to its role in cMYC recruitment to the chromatin, WDR5 became an attractive drug target in cMYC dependend cancers such as acute myeloid leukemia (AML). However, in WDR5 two distinct binding sites have been described: I) the WDR5-interacting site (WIN site) which binds e.g. to the SET-family methyltransferases.^15^ The WIN site has been successfully targeted by inhibitors such as OICR-9429^16^ or MM-598^17^. II) The WDR5-binding motif (WBM site) with a shallower binding surface responsible for cMYC recruitment has been also recently successfully targeted.^18^ Since blocking of the individual sites has not led to the anticipated therapeutic effects (probably due to not inhibiting all of the WDR5-mediated oncogenic functions)^19^, the development of PROTACs may offer a promising approach targeting all WDR5 scaffolding functions. So far, only WIN-site PROTACs have been published showing anticancer efficacy in AML cell lines.^20^

**Figure 1:**
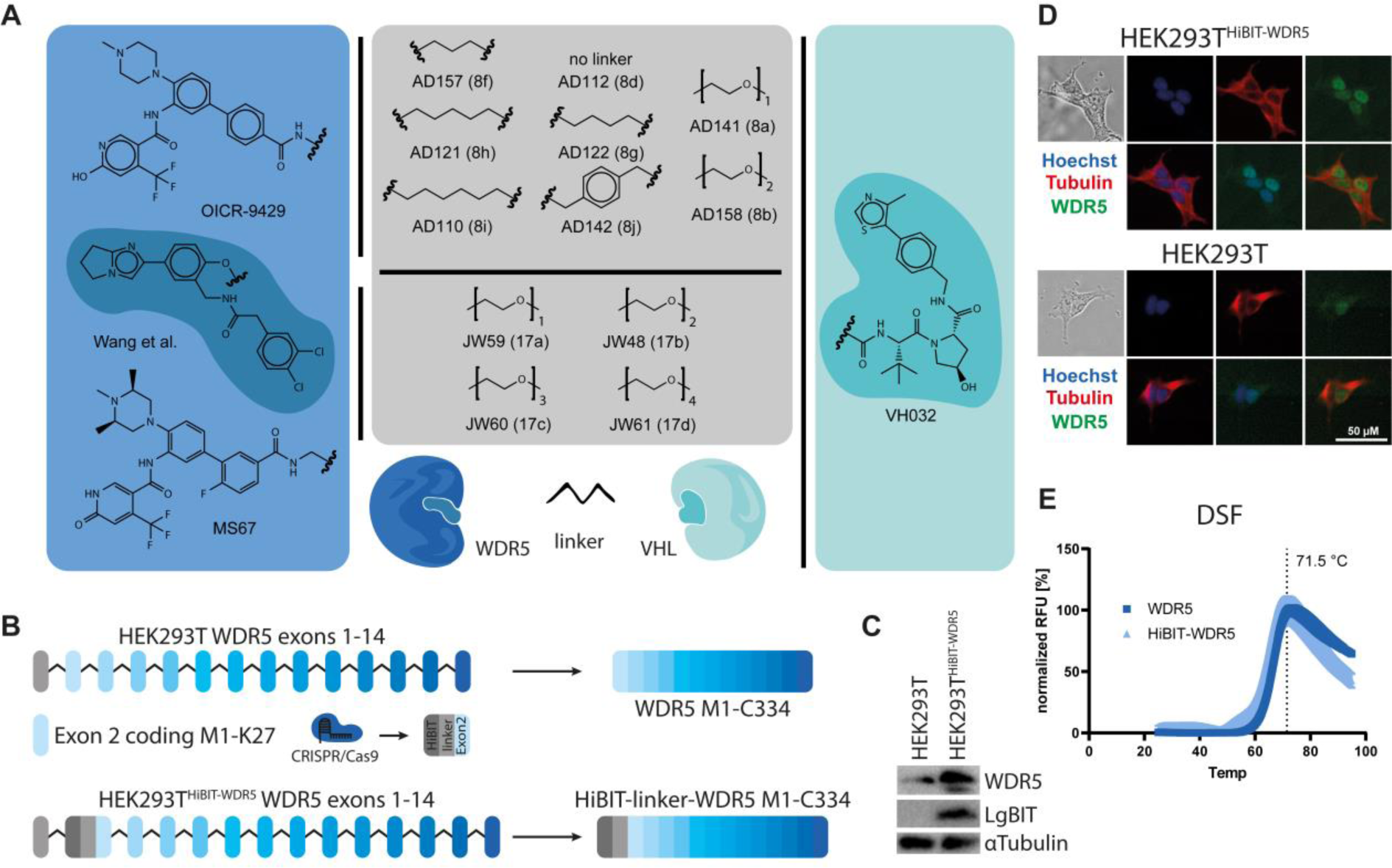
Compounds used in this study and the generation and validation of the tool cell line HEK293T^HiBIT-WDR5^. A) Left panel shows the different WDR5 ligands used for the synthesis. Middle panel depicts the different linkers that were used to link the target ligand to the E3 handle. Right panel displays VH032 as E3 handle which allows the recruitment of the PROTACs to VHL. B) Scheme of the genomic clustering of the WDR5 exons with Exon 2 encoding the start methionine (left panel). Final mRNA containing the HiBIT Tag at the N-terminus after splicing (right panel). C) In-cell ELISA for the determination of the cellular localization of unmodified and HiBIT-tagged WDR5. Hoechst staining (blue) is corresponding to the nucleus, while Tubulin staining (red) is corresponding to the cytosol. WDR5 (green) shows major localization in the nucleus. D) comparison of HEK293T wild type and HEK293T^HiBIT-WDR5^ cells through Western blot. WDR5 and Tubulin were detected through antibodies, while the LgBIT complementation of the HiBIT tag was measured via direct bioluminescence.

In this manuscript, we present a cell based assay pipeline for the comprehensive characterization of SAR (structure activity relationships) of PROTACs by monitoring critical steps of the degradation cascade. The basis of our study were three diverse chemical series of PROTACs with different levels of degradation activity. The developed assay cascade highlighted in particular the importance of stable and synergistic ternary complex formation in the cellular environment in order to achieve efficient PROTAC mediated degradation activity. Ternary complexes of one chemical series were further characterized by comparative structural biology studies. The established pipeline allows to track initial steps of the PROTAC mediated degradation process enabling a basis for the rational development of PROTACs and for identifying limiting steps in the degradation process.

## Results

### Generation of endogenously HiBIT tagged WDR5 as sensor for cellular protein levels

A key assay for PROTAC evaluation is the monitoring of endogenous cellular POI levels after PROTAC treatment. This analysis provides key data on PROTAC potency (DC_50_) as well as on the maximal level of POI depletion (D_max_). The most frequently used detection method of cellular protein levels is Western blotting. However, selective and well-validated antibodies are required and often the dynamic range of this method is insufficient and experiments are work intensive in particular if accurate determination of DC_50_ and D_max_ values is the goal. WB is particularly challenging for POIs with low cellular abundance. Additionally, the low throughput of Western blotting can be a limiting factor for the evaluation of large SAR series. To overcome these problems, sensory systems based on endogenous protein tagging by a fluorescent marker have gained popularity due to their large dynamic range and accuracy as well as the possibility of measuring detailed kinetic data on POI degradation. In this study we developed a HiBIT sensor cell line which allows detections of endogenous WDR5 protein levels based on the highly luminescent NanoLuciferase (NLuc). In this assay format, the NLuc was split into the HiBIT, an 11 amino acid long beta sheet and the remaining NLuc, called LgBIT. Without complementation, the NLuc is inactive whereas complementation leads to an active luciferase. For the generation of this sensor cell line, we used CRISPR/Cas9 technology in HEK293T cells with the goal to tag the N-terminus of WDR5 (Exon 2) with the small HiBIT tag (UniProt: P61964) as depicted in Figure 1.

Expression levels (mRNA) of relevant proteins including VHL, WDR5 and cMYC (SI Figure 4) and relevant proteins of the ubiquitin proteasomal system (UPS) (SI Table 1) were compared in HEK293T cells and a broad spectrum of leukemia cell lines. This analysis revealed no large differences in expression levels with only a slightly lower expression of WDR5 in HEK293T cells making this cell line a suitable host system to keep all assays in the same expression host and environment while containing all necessary markers of leukemia cancer cell lines. Following the CRISPR/Cas9 experiments and screening for luciferase signals as a selection marker, Sanger sequencing was carried out revealing only heterozygous insertion events. Despite the presence of two sequences peaks at the same locations, the HiBIT and linker sequence was present in the sequenced monoclone showing correct insertion (SI Figure 2). In order to rule out off-target LgBIT complementation caused by off-target HiBIT tagging, Western blots of the engineered cell line HEK293T^HiBIT-WDR5^ and the parent cell line HEK293T were carried out. Gratifyingly, we identified no additional bands apart from the anticipated WDR5-corresponding bands (Figure 1 C). N-terminal HiBIT-tagging of WDR5 increased the total amount of protein without a change in cellular localization and cellular morphology (Figure 1 D). Thus, the additional N-terminal amino acids may serve as a stabilizing tag increasing the cellular half-life as often observed when N-terminal sequences in a protein are altered. The influence of the HiBIT tag on the protein stability was tested using a heterologously expressed HiBIT-WDR5 fusion protein (as expressed in the HEK293T^HiBIT-WDR5^ cell line) and purified untagged WDR5 (His_6_-Tag cleaved). Stability was analyzed by ΔT_m_ assays (Differential Scanning Fluorimetry, DSF) for comparison which showed no thermal destabilization of WDR5 due to the HiBIT-linker fusion (Figure 1 E). However, we observed a second, not HiBIT tagged isoform in the Western blot analysis, which was apparent as a weak band. We concluded that since this isoform did not carry the HiBIT tag, it would not be detected in the complementation assays and the presence of this minor isoform did therefore not interfere with the resulting data. To exclude potential unspecific degradation effects on WDR5 caused by compound-induced toxicity, we assessed the viability of HEK293T^HiBIT-WDR5^ cells in presence of a representative set of WDR5 PROTACs or controls after 24 h. No significant decrease in cell viability was observed even at the highest PROTAC dose (7.5 µM) (SI Figure 7). Finally, we confirmed the cellular localization of the tagged WDR5 using in-cell ELISA in HEK293T and HEK293T^HiBIT-WDR5^ (Figure 1 D and SI Figure 5). These experiments confirmed the expected nuclear localization of HiBIT-WDR5 without observed changes in the cell morphology that could be potentially caused by the slightly higher expression levels and the HiBIT tag (Figure 1 C). Additionally, nuclear localization of WDR5 was validated by NanoBRET tracer titration in HEK293T^HiBIT-WDR5^ cells with a non-nucleus penetrating tracer molecule resulting as expected in non-detectable binding (SI Figure 6). Thus, the generation of the HEK293T^HiBIT-WDR5^ cell line granted a sensory system for monitoring live-cell protein levels. In addition, the established cell line allowed not only endpoint but also kinetic measurements on POI levels after PROTAC treatment.

**Figure 2:**
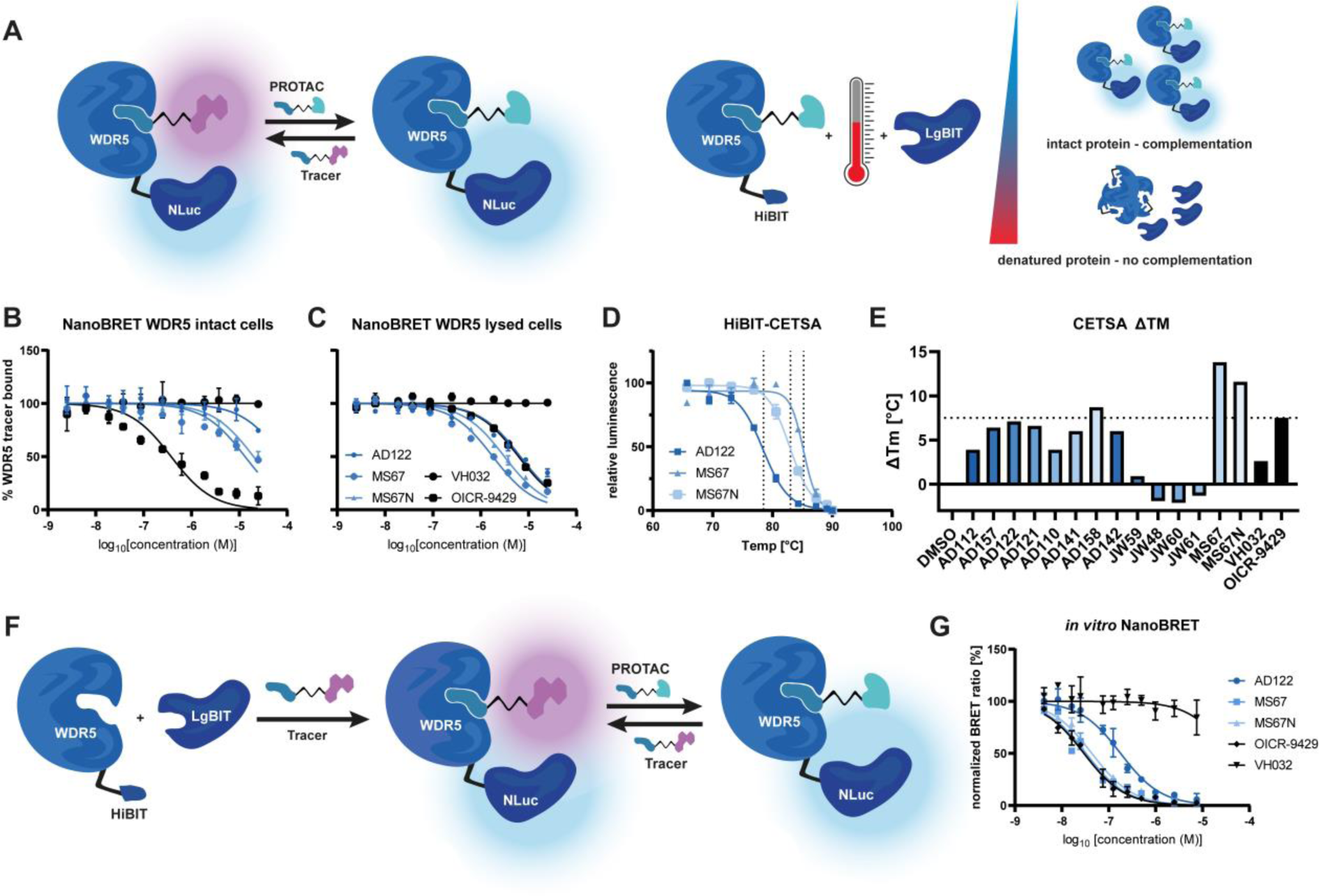
Binary complex measurements using NanoBRET and HiBIT technology. A) Left, graphical representation of the NanoBRET principle. BRET signal is observed with tracer-WDR5 binary complex. PROTAC binding leads to tracer displacement and loss of BRET signal allowing the estimation of % tracer bound to the protein and therefore measurement of PROTAC binary complex potency. Right, graphical representation of the HiBIT-based CETSA assay. HEK293T^HiBIT-WDR5^ cells are treated with compounds and exposed to a temperature gradient. After incubation, LgBIT complementation and cell lysis took place showing luciferase signal correlating with the amount of intact protein. Here, interaction of compounds did stabilize the protein leading to thermal shifts in the melting temperature. Intact (B) and lysed (C) cell target engagement of different PROTACs. As tracer molecule, the OICR-9429-based compound 19c from Dölle *et al.* was used.^21^ Displacement was measured for the different compounds after 2h incubation in presence of tracer and compound with (B) and without (C) digitonin for cell membrane disruption. Data was measured in biological replicates with error bars expressing the SD. n=2. D and E) HiBIT CETSA results as cellular melting curves (E) and thermal shifts expressed as bar diagram (E) Data was measured in biological replicates and expressed with error bars expressing the SD. F) Graphical representation of the *in vitro* NanoBRET assay. First, purified HiBIT-tagged WDR5 is complemented *in vitro* with the LgBIT protein to form a NanoLuc-WDR5 complex. This complex can subsequently be used to measure *in vitro* tracer displacement. G) exemplary data from *in vitro* NanoBRET measurements. n = 2.

**Figure 3:**
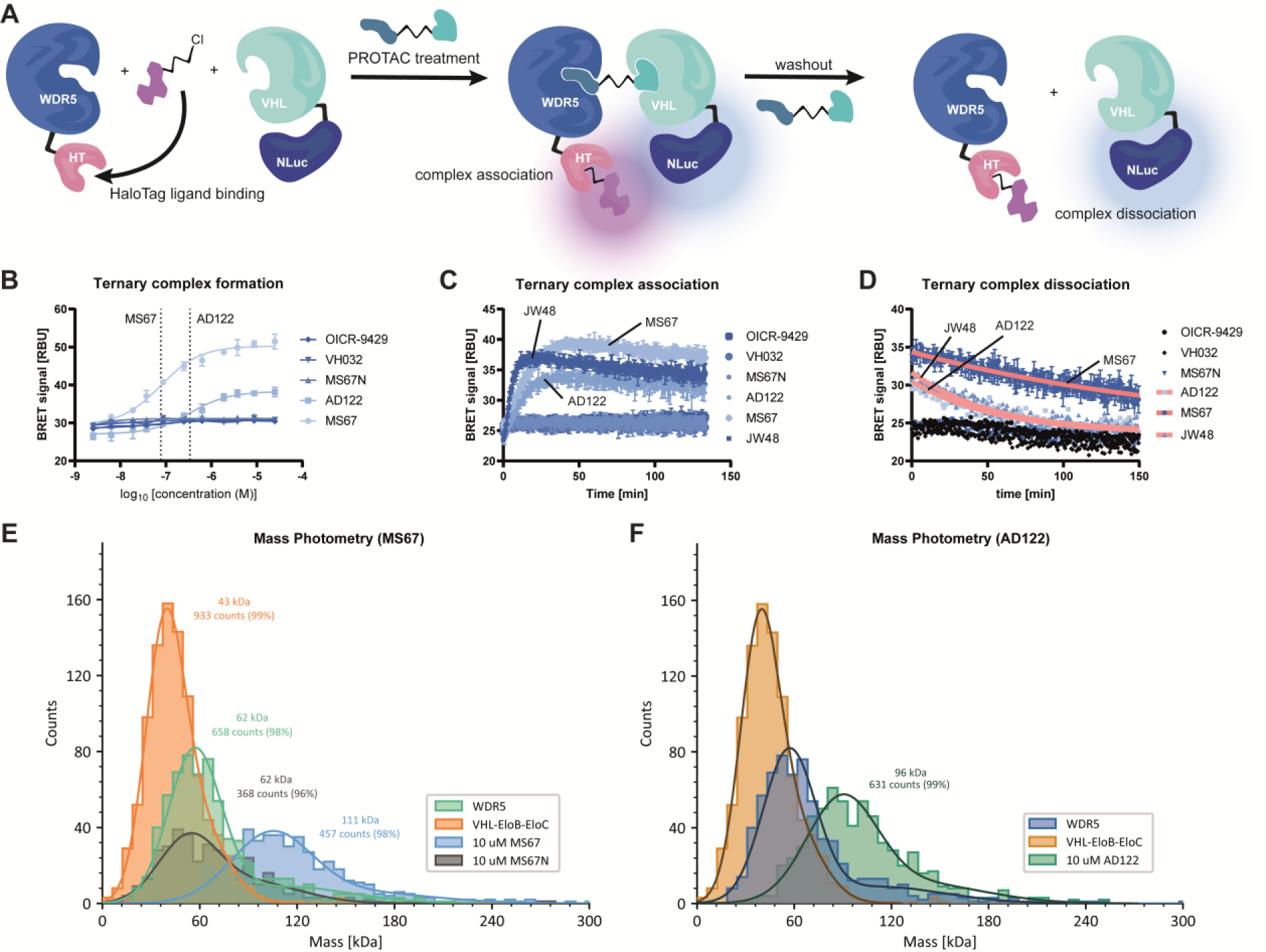
Ternary complex measurements using HiBIT and HaloTag technology technology (live cell) or mass photometry (*in vitro*). A) Graphical representation of the ternary complex formation assay. First, HaloTagged WDR5 and NLuc tagged VHL were overexpressed and treated with the HaloTag ligand for subsequent BRET measurements. After PROTAC treatment, the complex can form, leading to an increase in BRET signal through induced proximity between NLuc and HaloTag ligand. After measurement of association, the cells were washed to trigger dissociation of the complexes over the time. Fitting exponential decay curves yielded estimated half times, while the maximum during association delivered estimates association times. B) Ternary complex affinity determination using the established NLuc / HaloTag assay system. C) Ternary complex association kinetics in living cells. D) Ternary complex dissociation kinetics after PROTAC washout. Curves used for half-life calculation are shown in pink. All samples were carried out in biological replicates and error bars express the SD. n = 2 E) Histograms measured by mass photometry confirm strong ternary complex formation in the presence of MS67 but not using the inactive control MS67N. Recombinant proteins in the absence of PROTACs showed the expected molecular weight of the WDR5 as well as the VHL/ElonginB/C complex. F) Histograms measured by mass photometry confirm strong ternary complex formation in the presence of AD122. Additional experiment with other PROTACs and a MS67 concentration-response experiment are shown in supplemental figure 31.

**Figure 4:**
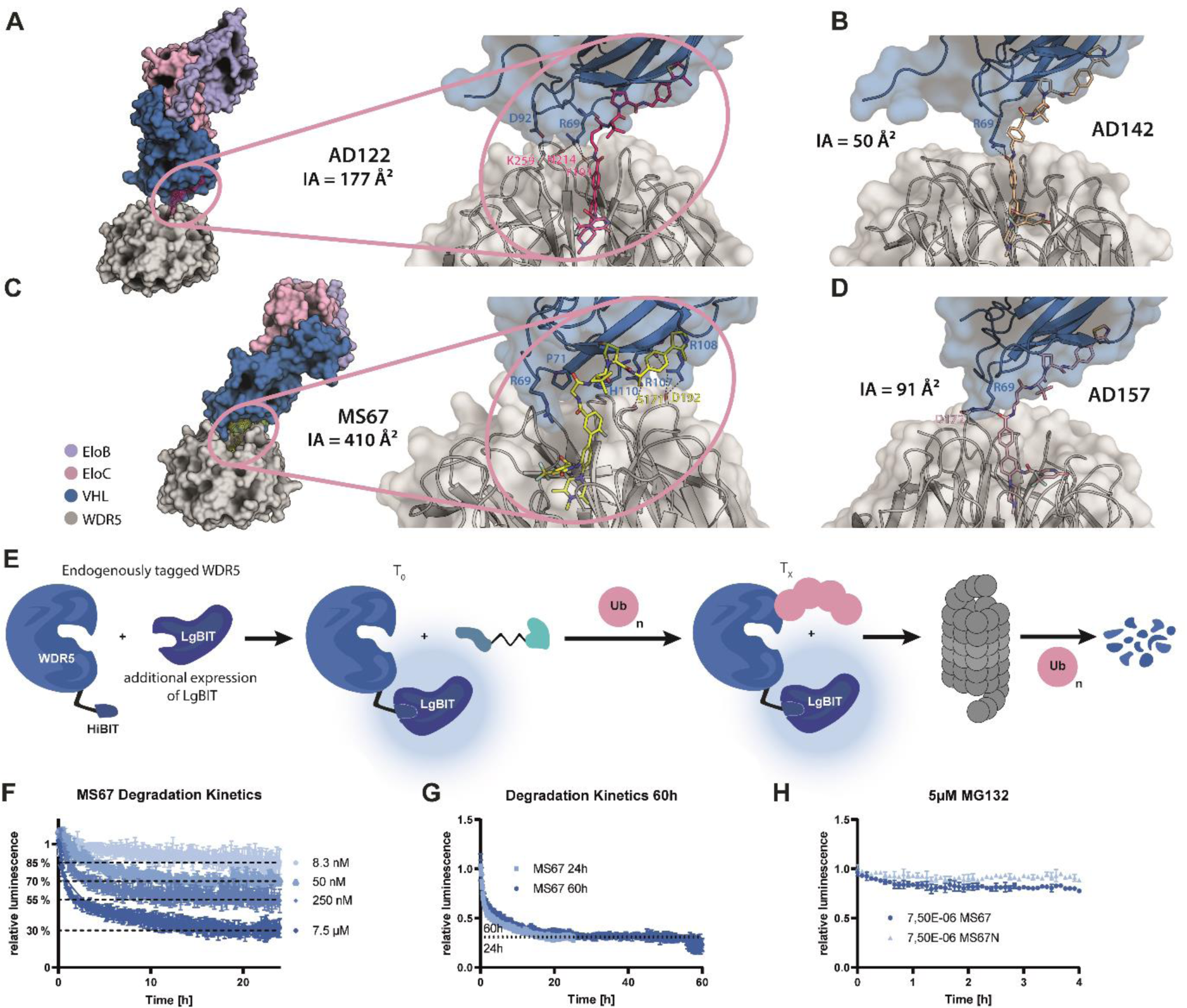
X-ray structures of ternary complexes and degradation measurements of the PROTAC-induced degradation of WDR5 in live cells and lysed HEK293T^HiBIT-WDR5^ cells. A) overall structure of VHL-EloB-EloC, WDR5 and AD122 (left panel) and detailed binding interface (right panel) with indicated interface area (IA) (pdb: 7q2j). B) detailed binding interface of AD142 (pdb: 8bb5). C) overall structure of VHL-EloB-EloC, WDR5 and MS67 (left panel) and detailed binding interface (right panel) (pdb: 7jtp). D) detailed binding interface of AD157 (pdb: 8bb4). E) Scheme of the assay system used for the measurements. The HiBIT-tagged WDR5 expressing cell-line was transfected with the LgBIT expression vector for NLuc complementation. After proteasomal degradation, the loss of NLuc signal was detected as correlation to the protein depletion. F) Degradation kinetics of different doses of MS67 with the respective D_max_ values (dotted lines) after 24h. G) 60h kinetic measurements using 7.5 µM MS67 showing a lack of recovery after 24h and a correlation between the 24h D_max_ and the 60h D_max_. H) Degradation kinetics of 7.5 µM MS67 and MS67N after 5µM MG132 treatment. IAs were calculated using PISA.^27^

**Figure 5:**
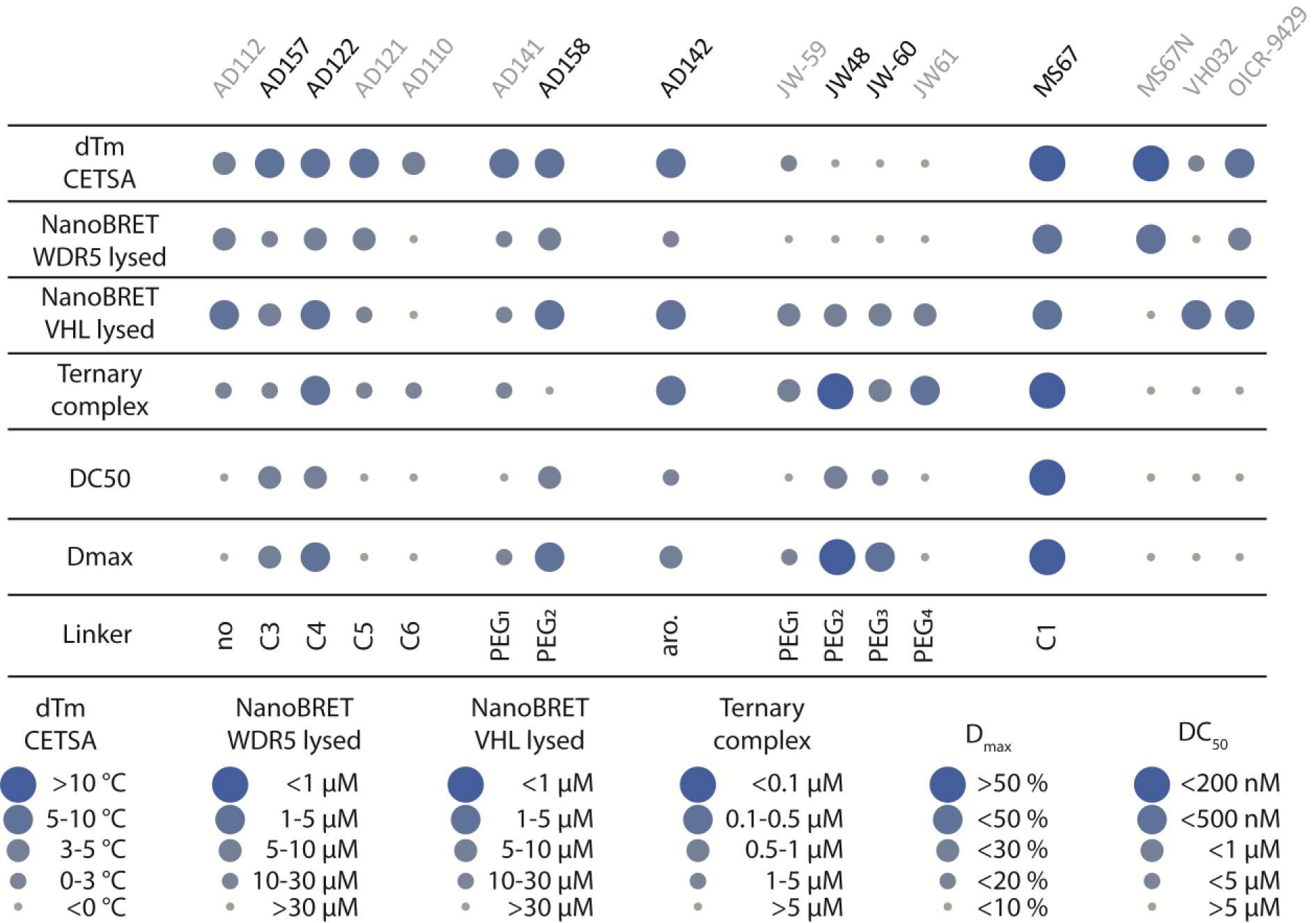
Dot plot illustration of the obtained data in all characterization assays. The measured potencies and temperature shifts are represented by spheres of varying size and color as indicated in the figure caption.

**Table 1:** Affinity data measured for binary complex formation. Published DSF data was taken from Dölle et al.^21^ DSF and CETSA data was expressed in °C. CETSA was measured in biological replicates. The melting temperature was determined by graphing the data points and determining the IC_50_. IC50 values for NanoBRET were taken from biological replicates and were expressed in µM. Values higher than 30 µM are shown as >30 µM due to increasing error for values > 30. n=2 For the in vitro NanoBRET data, values were taken as IC_50_ values from duplicate measurements. n=2

| Compound | Published<sup>1</sup><br>*in vitro*<br>DSF [°C] | CETSA<br>thermal<br>shifts<br>[°C] | NanoBRET<br>WDR5<br>[μM] | Lysed<br>NanoBRET<br>WDR5<br>[μM] | NanoBRET<br>VHL [μM] | Lysed<br>NanoBRET<br>VHL [μM] | In vitro<br>NanoBRET<br>[nM] |
| --- | --- | --- | --- | --- | --- | --- | --- |
| AD112 | 15.6 | 3.9 | >30 | 8.3 | >30 | 1.9 | 279.2 |
| AD157 | 10.4 | 6.4 | >30 | 15.2 | >30 | 5.2 | 131.6 |
| AD122 | 13.2 | 7.1 | >30 | 6.7 | >30 | 1.9 | 178.8 |
| AD121 | 9.7 | 6.6 | >30 | 6.7 | >30 | 8.2 | 92.0 |
| AD110 | 3.5 | 3.9 | >30 | >30 | >30 | >30 | 515.3 |
| AD141 | 15.3 | 6 | >30 | 11.2 | >30 | 5.1 | 189.7 |
| AD158 | 7.7 | 8.7 | 8.2 | 5.6 | 15.2 | 2.2 | 62.1 |
| AD142 | 17 | 6.1 | >30 | 18.4 | >30 | 16.3 | 180.1 |
| JW-59 | 7 | 0.9 | >30 | >30 | >30 | 26.0 | >3000 |
| JW48 | 5.2 | -1.9 | >30 | >30 | >30 | 3.5 | >3000 |
| JW-60 | 3.6 | -2.1 | >30 | >30 | >30 | >30 | >3000 |
| JW61 | 3.4 | -1.3 | >30 | >30 | >30 | 1.7 | 1780 |
| MS67 | n.d. | 13.8 | 13.2 | 1.8 | >30 | 6.8 | 32.6 |
| MS67N | n.d. | 11.6 | 19.9 | 3.1 | >30 | 5.4 | 43.6 |
| VH032 | n.d. | 2.6 | >30 | >30 | 1.8 | 4.3 | >3000 |
| OICR-9429 | 13.2 | 7.5 | 0.4 | 6.2 | >30 | 3.3 | 29.6 |

### Detection of binary complexes

Binary complexes of the PROTAC with the POI as well as the E3 ligase are crucial for subsequent protein degradation. Sensor systems monitoring binary complex formation provide “on-target” assays and important binding data (K_D_ values) that monitor if a PROTAC efficiently reaches the POI. Interference of PROTAC binding to the POI or E3 might be due to non-optimal installation of linker moieties resulting in weak interactions compared to the parent ligand. However, strong POI engagement may also result in an early onset of the so called hook effect in which the binary complexes compete with the formation of ternary complexes leading to abrogation of POI degradation. In order to monitor binary complex formation, we used cell (NanoBRET) based assays which served not only assessing target engagement of the PROTACs, but also membrane penetration of PROTACs by comparing apparent K_D_ values measured in intact and permeabilized cells as previously reported for one of the developed PROTAC series.^21^

First, we assessed cellular target engagement of all compounds towards WDR5, using the most suitable tracer identified by Doelle and colleagues (19c).^21^ Subsequently, PROTAC titrations revealed weak cellular compound-WDR5 potency of often > 30 µM in intact cells, whereas parent compounds showed potent binding in live cells (Table 1 and Figure 2 A-C). Binary complex formation of the compounds towards VHL resulted in comparable potencies (Table 1). However, in permeabilized cells (digitonin-treated) most PROTACs revealed *IC*_50_ values of < 10 µM suggesting that limited cell penetration was the most likely reason for weak binary complex formation in living cells. (Table 1).

To verify these data, we established a cellular thermal shift assay (HiBIT-CETSA) using the sensitive HiBIT-tagged WDR5 (HEK293T^HiBIT-WDR5^) as a detection system.^22^ Surprisingly, a relatively high cellular melting point of WDR5 was determined (71.4 °C) while PROTAC addition further stabilized WDR5 towards temperature-induced denaturation. We obtained relative shifts in melting temperature up to 13.8 °C for MS67 increasing the WDR5 melting point to 85.2 °C (Table 1 and Figure 2 D). The measured compounds clustered into three groups. First, the OICR-9429 based series (AD) showed thermal shifts comparable to the parent compound (7.5 °C). However, AD110 and AD112 revealed decreased thermal shifts indicating weakened affinity for WDR5 possibly due to unfavorable arrangement of the bivalent compound (Figure 2E). Second, MS67 and MS67N which used a more optimized WDR5 ligand showed as expected the highest ΔTm shifts. Surprisingly, the JW PROTAC series which contained a structurally diverse WDR5 ligand did not form detectable binary complexes with WDR5. This surprising result was confirmed by the lack of stabilization of WDR5 in the CETSA assay as well as low affinities in cellular NanoBRET target engagement assays, while the other series formed complexes in living cells with micromolar affinities. (Table 1). Comparison of ΔTm shifts in intact and lysed cells resulted in identical values suggesting that cells were lysed prior to reaching the high melting temperature of WDR5. However, the CETSA experiments provided binding data for PROTACs for which we were not able to measure target engagement by NanoBRET using endogenous tagged WDR5.

To validate the binary complex assays, HiBIT-tagged WDR5 protein was complemented *in vitro* with the LgBIT protein (Figure 2 F). The reconstituted NLuc-WDR5 protein was used in NanoBRET assays revealing a tracer K_D,app_ of 30 nM (SI Figure 29). Using this value as tracer concentration in the subsequent compound titrations, we were able to successfully establish an *in vitro* NanoBRET assay system for WDR5 demonstrating that the N-terminal HiBIT-linker attachment to WDR5 did not alter protein stability and function, but it also provided valuable binding data for the characterization of WDR5 PROTACs (Figure 2 G). In this assay, OICR-9429 did outcompete all other PROTACs revealing an affinity of ∼ 30 nM. The PROTAC affinity for WDR5 show excellent correlation with CETSA data and revealed that the warheads used in the JW series (> 1 µM) formed only weak binary complexes compared to the other PROTAC series (Table 1).

Assessment of the binary complexes provided interesting insights into the mechanism of the tested PROTACs. Regardless the sub 10 µM binding affinity in lysed cells towards VHL and WDR5, affinities in intact cells were significantly lower possibly due to poor membrane penetration of the used PROTACs. Interestingly, despite all parent compounds have been published to strongly bind WDR5, the JW series did not form detectable binary complexes in cells, but they showed weak interaction in lysed cells. The weak interaction of these PROTACs was intriguing based on the published activity of these PROTACs as potent WDR5 degraders.^21^

### Ternary complex formation in live cells are key events for successful POI degradation

PROTAC induced ternary complex formation of the E3 ligase and the POI has been considered as a key event and possibly a rate determining step of PROTAC mediated degradation. Synergy in the formation of the ternary complex has been described as an important requirement for efficient POI degradation ^23^ suggesting that the formation of synergistic POI/PROTAC/E3 complexes may depend on the compatibility of E3/POI interface as well as the orientation induced by the PROTAC linker. Optimization of the PROTAC linker, including rigidifying it has been reported as a strategy improving PROTAC efficacy after the optimal linker length and attachment point has been identified.^24^ However, also ternary complexes that assemble none-synergistically may lead to efficient POI degradation.^25^ Importantly, binary complex formation competes at high PROTAC concentrations with the formation of ternary complexes resulting in loss of POI degradation, a property that has been described as the hook effect.^26^ To date, PROTAC induced ternary complex formation has been mainly studied by biophysical methods such as ITC, fluorescent based assays and other methods that require recombinant proteins. Due to the limited throughput of these work intensive methodologies, studies that compare diverse series of PROTACs as well as PROTACs that form binary complexes but that do not result in POI degradation have not been published so far. To this end, we developed a luminescence sensor system based on a BRET pair consisting of a HaloTagged WDR5 construct together with a NLuc-tagged VHL sensor. The combination of these two sensors established an assay with which the formation as well as dissociation of the ternary WDR5-VHL-PROTAC complex can be monitored in cellular environments at a throughput that allows the assessment of all available WDR5 degraders as well as poorly or inactive PROTAC molecules (Figure 3 A).

The established assay allowed the monitoring of ternary complex formation resulting in the determination of apparent ternary complex *K*_D_ values in cells (Figure 3 B) as well as assessment of ternary complex association and dissociation kinetics (Figure 3 C and D). While representative graphs are shown in Figure 3, data for all PROTACs have been compiled in Table 2. Interestingly, the signal intensity was observed to be highly dependent on the measured PROTACs, a property which is likely caused by differences in the orientation and distance of the sensors in PROTAC induced ternary complexes. As expected, the relative BRET signal did not correlate with the association strength of ternary complexes but it was merely a function of the orientation and distance of BRET sensors. For instance, within the AD PROTAC series, the comparison of the maximum detected BRET signal correlated with the linker length e.g. for the aliphatic linkers AD110>AD121>AD122>AD157 where the BRET intensity decreased with increasing linker length and therefore WDR5-VHL distance. The affinity determination yielded apparent association constants for all PROTACs except for AD158 (and negative controls). The relative affinities did span several order of magnitudes including PROTACs that induced a highly cooperative complex formation such as MS67 (apparent K_D_: 60 nM) to moderate affinities induced for instance by AD112 (3 µM) (Table 2). Furthermore, comparing BRET data across the three chemical series we realized that information on complex conformations can be estimated from the BRET signal strength (SI Figure 16). It is interesting to note that MS67, which had a unique ternary complex arrangement resulted in the strongest BRET signal. The AD series gave rise to a lower BRET signal, in agreement with the significantly different orientation of the WDR5-VHL/elonginB/C arrangement. Finally, the JW series showed even a smaller maximum BRET signal, indicating a potential third conformation which we were however unable to elucidate by structure studies due to difficulties crystallizing the ternary complex of this series.

**Table 2:** Affinity measured for the ternary complexes together with the estimated association times and half-life for the complexes. Affinity data was measured in biological replicates with each technical duplicates while kinetics were measured in biological replicates. n=2

| PROTAC | Ternary complex<br>affinity [nM] | Estimated ternary complex<br>association time [min] | Ternary complex half-life [min] |
| --- | --- | --- | --- |
| AD112 | 3092.5 ± 1568.0 | 40 | 107 |
| AD157 | 3234.0 ± 41.0 | 19 | 53 |
| AD122 | 394.5 ± 56.5 | 36 | 40 |
| AD121 | 2552.0 ± 605.0 | 20 | >150 |
| AD110 | 3173.0 ± 1457.0 | 10 | 61 |
| AD141 | 3719.0 ± 189.0 | 50 | >150 |
| AD158 | - | - | - |
| AD142 | 207.3 ± 49.9 | 50 | 52 |
| JW-59 | 565.1 ± 43.0 | 8 | >150 |
| JW48 | 95.0 ± 22.2 | 17 | 36 |
| JW-60 | 766.3 ± 117.1 | 9 | >150 |
| JW61 | 424.1 ± 24.2 | 8 | 47 |
| MS67 | 63.8 ± 13.7 | 45 | 134 |
| MS67N | - | - | >150 |
| VH032 | - | - | >150 |
| OICR-9429 | - | - | >150 |

Gratifyingly, the developed assay system allowed also investigation of the association and dissociation kinetics of ternary complexes which provided further insights into the biophysical characteristics of the investigated PROTAC induced ternary complexes. While the overall time courses for complex formation were between approximately 10 and 50 minutes, PROTACs based on the JW series induced significantly faster complex formation compared to the OICR-9429-based analogues. Compared to the complex association times, dissociation of the complexes was drastically slower. Here, complex half-lives ranged from 36 minutes for JW48 indicating fast on- and off-rates, to 134 minutes for MS67 also correlating with the respective slow on-rate of 45 minutes of this PROTAC. These findings showed that similar PROTAC induced-ternary complex affinities may have significantly different association and dissociation kinetics. As an additional *in vitro* validation for successful ternary complex formation, all PROTACs which were used for structure determination as well as the negative control compound MS67N were studied by mass photometry (Figure 3 E and F). In this experiment the molecular weight of the recombinant proteins was accurately determined. Incubation with MS67 and AD122, but not the negative control MS67N resulted in a mass shift expected for the ternary complex. Formation of ternary complexes was also confirmed for AD157, AD142 and JW48 and concentration-response experiments revealed a shift towards the expected mass of the ternary complex with increasing PROTAC concentration (SI Figure 31).

After characterizing the assembly and dissociation of ternary complexes in live cells, we were interested in the molecular mechanisms that may determine ternary complex stability. To address this question, we successfully determined the crystal structures of the VHL-EloB-EloC-PROTAC-WDR5 complex with bound AD122, AD157 and AD142 (Figure 4 A-D and SI Figure 25). The complex with MS67 has already been published previously^20^. All structures were refined at a resolution range between 2.2-2.8 Å and ligands as well as interacting side chains were well-defined by electron density (SI Table 6 and SI Figure 25). As illustrated in Figure 4, two major overall arrangements were observed (panel A and C). Albeit PROTACs from the AD series showed slightly altered interactions resulting in different interface areas (IA), all complexes displayed a relatively small interface similar to the previous published complex of MS33^20^ (PDB: 7JTP) (Figure 4 A, B and D). In contrast to the AD-series, the ternary complex induced by MS67 adopted a much more extensive protein-protein interface (Figure 4 C) in which WDR5 was shifted approx. 25 Å and rotated approx. 90° (SI Figure 26). In good correlation with the measured D_max_ values, the most potent degrader MS67 displayed the largest calculated protein-protein interface (without PROTAC) area of approx. 410 Å^27^, while AD122, in agreement with its potency as a degrader, had with 177 Å^2^ the largest IA among the AD series. The most prominent interactions between VHL and its AD122-induced neosubstrate WDR5 were formed by hydrogen bonds between R69^VHL^ and the backbone of N214^WDR5^ and Y195^WDR5^. Additionally, D92^VHL^ and K259 ^WDR5^ formed hydrogen bonds which were not identified in other AD series structures. Compared to AD122, both AD157 and AD142 showed a less pronounced interface of approx. 91 Å^27^ and 50 Å^27^, respectively. Here, the only polar interaction between VHL and WDR5 was formed by R69^VHL^ for AD157 while R69^VHL^ in the AD142 structure only displayed interaction towards the bound ligand (Figure 4 B and D. Both PROTACs had similar degrading efficiency (Table 3). However, it is noteworthy that the interactions between WDR5 and VHL were rather weak and according to PISA^27^ these interfaces are not energetically favored with calculated ΔG close to 0 kcal/mol. Interestingly, the interface between the PROTACs and the respective proteins were much more extensive compared to the WDR5-VHL protein interfaces. AD122 displayed a calculated interface of approx. 500 Å^27^ with WDR5 and additional 400 Å^27^ with VHL, yielding a total interface of ∼ 900 Å^27^. In accordance with the measured affinities for the induced ternary complexes ΔG of these interactions were strongly negative with −4.3 kcal/mol for WDR5 and −2.9 kcal/mol for VHL, respectively. Thus, the combined interfaces of the PROTAC with both binding sites are most likely the main driving force for complex formation. For the whole AD series, all PROTACs exhibit similar calculated values for the interface areas and ΔG, while MS67 has an even higher combined calculated interface of over ∼1000 Å^27^ (Figure 4 C). However, the increased interface of MS67 was mainly due to three modifications on the original WDR5 scaffold and one on the VHL-ligand (Figure 1 A). These modifications have been introduced to improve affinity to both proteins, which resulted in stronger binary as well as ternary complexes and an increased overall surface area^27,28^.

**Table 3:** DC_50_ values for 4h, for the AUC of 24h and the D_max_ between 23 and 24h. D_max_ was calculated as mean from the last 12 measurements from both duplicates with SD calculated.

| Compound | DC<sub>50</sub> (4h) in nM | DC<sub>50</sub> (AUC) in nM | D<sub>max</sub> (23-24h) at 7.5 μM in % | Degradation rate (7.5 μM) in min<sup>-1</sup> |
| --- | --- | --- | --- | --- |
| AD157 | 722 | 1316 | 21 ± 11 | 0.48 |
| AD122 | 896 | 767 | 38 ± 11 | 0.39 |
| AD158 | 876 | 1244 | 32 ± 7 | 1.07 |
| AD142 | 1557 | 215 | 28 ± 11 | 1.04 |
| JW48 | 551 | 4275 | 55 ± 6 | 0.75 |
| JW-60 | 1483 | 2675 | 41 ± 5 | 1.3 |
| MS67 | 148 | 209 | 69 ± 4 | 1.22 |

Although these calculated interaction surfaces must be treated with caution, there seemed to be a good correlation between the interaction surface and ternary complex affinity. However, crystal packing may additionally influence domain orientations. We were unable to crystallize a ternary complex of the JW series, probably due to the weak binary complex affinity. However, attempts yielded crystal structure of a binary complex of VHL and a partially resolved JW48 (SI Figure 27). Comparison with the structure of the parent compound suggest that the exit vector might be similar to the one observed in the MS67 complex, suggesting that also for these PROTACs a favorable ternary complex interface can be formed.

The successful determination of the ternary complex formation in cells provided key parameters of the central ternary complex formation and stability and insights into the degradation mechanism of the tested PROTAC SAR series. While all negative controls were as expected unable to form ternary complexes, we were able to measure even weak ternary complexes with the established assay. Surprisingly, PROTACs with undetectable binary complexes in intact cells were validated as proximity inducers as well as efficient degraders of WDR5, highlighting the importance of testing the formation of ternary complexes in cellular environments as a key parameter for PROTAC optimization.

### Proteasomal degradation of WDR5 in live cells

Target protein degradation is the ultimate endpoint in the multistep process of PROTAC-induced protein degradation before the PROTAC is recycled for another degradation cycle. Typically, the decrease of total protein is measured in cell lysates (after lysis e.g. WB) possibly with subsequent trypsin digest of the total proteome for protein abundance measurements (proteomics). Since the targeted protein degradation process is highly dynamic, cell lysis attenuates the degradation cascade resulting in a loss of information on time dependent degradation processes. The least invasive method for live cell detection of endogenous POI concentration, is represented by the HiBIT-tagged cell line (HEK293T^HiBIT-WDR5^) which harbored only an addition short sequence inserted in the N-terminal coding sequence of the endogenous locus. Live tracking of this cell line allowed time-dependent measurements of WDR5 abundance after PROTAC treatment (Figure 4 E). This data was subsequently used for determination of the key parameter DC_50_ and D_max_, yielding the standard parameters for PROTAC characterization at different time points and PROTAC concentrations.

To assess live cell WDR5 protein levels after PROTAC treatment, cells were treated with PROTACs in dose-response using live tracking of WDR5 over 24 h. Assay quality was highly dependent on the PROTAC efficiency (D_max_) and potency (DC_50_). MS67 as the most efficient WDR5 degrader, provided high quality degradation kinetic data. Dose response experiments revealed a highly concentration dependent D_max_ of around 70% (indicated by the dotted lines in Figure 4 F) at the highest concentration used (7.5 µM) without showing signs of a hook-effect, indicating that the concentration of 7.5 µM was insufficient to outcompete ternary complex formation by binary complexes. Maximum level of degradation was stable up to 60h (Figure 4 G). The dependency of WDR5 degradation on the proteasomal system was confirmed by co-treatment with the proteasome inhibitor MG132 during kinetic WDR5 degradation measurement. As expected, this experiment revealed attenuation of degradation, confirming a proteasome dependent mechanism (Figure 4 H).

Next, for comparison of the initial degradation rates and therefore the speed of equilibrium adjustment, the degradation rate λ was calculated (table 3). After determination of the degradation rates, DC_50_ values were extracted from the kinetic degradation curves in time-dependent manner. Interestingly, after 2h of PROTAC treatment, the DC_50_ did not show any change in potency indicating that the initial degradation phase was shorter than 2h resulting in an equilibrium state as observed for MS67. Calculation of the DC_50_ of all studied PROTACs revealed sub-micromolar degradation potencies for nearly all tested compounds which was significantly lower than the estimated binary complex affinity (table 3). Next D_max_ parameters were extracted from the measured kinetic data. Apart from MS67 and JW48, none of the PROTACs revealed higher D_max_ values than 50%. Since both of these values were proven to be time dependent, analysis of the entire time courses of the monitored degradation kinetic by area under the curve (AUC) determination resulted in effective degradation concentrations for 24h of degradation duration, giving a single DC_50_ value containing all kinetic parameters such as initial degradation rate and time at D_max_ (table 3). The generated data suggested that kinetic measurement of live cell protein levels using our WDR5 sensor cell line can yield a comprehensive characterization of equilibrium and kinetic degradation parameters.

## Discussion

Rational development of PROTACs has been hampered due to the lack of available assay systems that would identify rate limiting steps of the PROTAC induced degradation process. Currently, Western blotting has evolved as the main screening tool for the identification of active PROTACs within a chemical series of structural variants which may comprise diverse linker attachment points, linker length and chemistry as well as E3 ligands. However, as the optimal time point and concentration may vary considerably between different PROTACs, and Western blotting lacks throughput and as an end point assay the possibility of continuous monitoring of protein degradation, the activity of a PROTAC may be misjudged. Importantly, the rate limiting step of PROTAC efficacy in the degradation cascade remains elusive as only the endpoint of protein degradation is measured.

In this study we developed fluorescent based sensors assessing key steps of the PROTAC induced degradation cascade such as membrane penetration, the formation of binary complexes with the POI and the E3 ligase as well as the kinetics and the relative affinity of PROTAC induced ternary complex formation and its final dissociation. The developed assays allowed the observation and assessment of these early and critical steps of the degradation cascade in living cells and they had sufficient throughput to evaluated a series of diverse PROTACs developed in our laboratories targeting WDR5. To study the effect of the different PROTACs on the multistep process of WDR5 degradation by the ubiquitin system, we generated a sensor cell line for live cell WDR5 level tracking. Due to the small size of the HiBIT tag (11 amino acids), we expected the least impact on WDR5 function. Indeed, we confirmed that apart from a moderate increase in cellular WDR5 protein levels potentially due to a stability increase based on the N-terminal HiBIT fusion, the protein was degraded similarly to wild type WDR5.^21^

In our kinetic characterization of the degrader set, we discovered that none of the investigated WDR5 degrader was able to completely degrade WDR5. However, the level of degradation may be cell line dependent. Previously we reported, that low VHL levels might be limiting WDR5 degradation.^21^ Additional explanations for the lack of complete POI degradation might be that 1) part of the WDR5 protein is protected by chromatin as described for Aurora A PROTACs^29^, 2) insufficient nuclear penetration of the PROTACs and 3) a fast WDR5 re-synthesis rate resulting in an equilibrium between re-synthesis and PROTAC induced degradation.

Target engagement assays (POI and E3 ligase) displayed only weak interaction of the PROTAC in intact HiBIT cells. Comparison with the same measurements in lysed cells revealed that cell penetration was the main reason for the observed low target engagement. In order to confirm the target engagement with HiBIT-WDR5 by a second biophysical technique, we developed a HiBIT CETSA assay that showed robust shifts for most PROTACs. However, the high melting temperature of WDR5 lysed the cells before a melting transition was observed resulting in identical data in lysed as well as intact cells. The most active degrader MS67 displayed a NanoBRET EC_50_ value of 13.2 μM for WDR5 in intact cells that was reduced to 1.8 μM in lysed cells and a melting temperature increase of 15.8 °C in the CETSA assay (Table 1). The most efficient degrader of the AD series showed binary complex formation only detectable in the lysed mode with EC_50_ values of 6.7 μM and 1.9 μM for WDR5 and VHL, respectively. However, in transient transfections assays published during the development of this series by our group, the interaction of these PROTACs had been demonstrated probably due to higher protein concentration and potentially due to non-nuclear localization of the ectopic WDR5 protein.^21^ Interestingly, none of the JW series PROTACs showed detectable target engagement towards WDR5 in live cells but affinity in the low μM regions were detected for the E3 ligase VHL. Intriguingly, none of the JW series PROTACs revealed a melting temperature shift in the CETSA assay despite the good WDR5 degradation activity of JW48. Thus, using only binary complex formation, the JW series may have been discarded by an initial SAR analysis based only on this property.

The most intriguing observation was the strong correlation of efficient WDR5 degradation and the strength of ternary complex formation (Figure 5). This is most dramatically shown for the degrader JW48 which displayed no detectable binary complex formation in intact cells but formation of a strong ternary complex (apparent K_D_: 95.0 ± 22.2 nM). As expected, also the most potent degrader MS67 exhibited potent formation of ternary complexes (apparent K_D_: 63.8 ± 13.7 nM). Next to this tight interaction, MS67 was able to induce a different interaction interface possibly explaining the superior behavior of MS67 compared to the other PROTACs, which showed a weaker interaction face due to the tilted orientation of the VHL-EloB-EloC complex. The only exception was AD158, which revealed an undetectable ternary complex but a modest DC_50_ (876 nM after 4h) and D_max_ (32 ± 7%). The combined data are illustrated in figure 4. Thus, from our analysis we concluded that a combination of assays reporting ternary complex formation together with the kinetic degradation assays represent the most suitable assay parameters for SAR analysis during PROTAC development.

## Author contributions

Manuscript was drafted and figures were designed by MS and edited by KS, AK and SK. Conceptualization, experimental design, cloning, protein purification, establishment of assays and was carried out by MS. Protein purification for X-ray crystallography as well as structure determination was carried out by AK. Compounds were provided by AD, JW, XY and JJ. Scientific supervision by SK and JJ.

## Acknowledgements

Special thanks to Benedict-Tilman Berger (BTB) Mohit Misra (MM) and Yves Matthes (YM). All were very helpful in discussing either cellular assays (BTB), mass photometry (MM) or CRISPR/Cas9 related questions (YM). Furthermore, additional thanks to YM and the Ciulli laboratory for providing pX330 and the expression plasmids.

## STAR Methods

### Generation of WDR5 HiBIT-tagged HEK293T cell line using CRISPR/Cas9

For CRISPR/Cas9 experiments, the gRNA needed to be cloned into the pX330 plasmid harboring the Cas9 gene. For this, gRNA was designed to introduce a double strand break at the N-terminus of the canonical isoform of WDR5, which starts at Exon 2. This gRNA (CACCGGCUGGCCGCACAGGAGACAAGUUU) was ordered in primer strands containing sticky ends for ligation after *Bbs*I digestion of pX330 after annealing of both primers (fw: CACCGGCTGGCCGCACAGGAGACAA; rv: AAACTTGTCTCCTGTGCGGCCAGCC). The ligated plasmid (pX330-WDR5) was checked by sanger sequencing. For the introduction of the HiBIT tag, a parent plasmid was cloned, harboring the HiBIT sequence together with a linker (NRIRGSSGGSS) and specific primers were designed containing 90 nt homology arms into the endogenous locus of WDR5 Exon 2 to PCR amplify the homology directed repair (HDR) template (fw: GGCTCCCTGTTCTGCATCTCGCTCAACAGACTGCCTCTGTCACCGGGTCCCTCCAGCCTTGTCTCCTGTGCGGCCAGCGTCAGAGCCATGGTGAGCGGCTGGCGGC; rv: CTGGGCACGCACCTTGCTCTGAGTGGCGGATGACGAAGGGGTTGGCTGTGCTCTGGCGGCCTCGGTCTCGGGCTTCTTCTCCTCCGTCGCGCTGCTGCCGCCGC). HEK293T cells were used in passage 9 for transfection with equal amounts (400 ng) of pX330-WDR5 and HDR template were transfected using FuGENE HD following the manufactureŕs recommendation. The cells were passaged for 10 days and single cell dilutions were carried out on 384-well clear bottom plates. After 8 days, single colonies formed and the plates were split into two copies, where one copy was kept in culture while the second was screened for luminescence, using the Nano-Glo® HiBiT Lytic Detection System (PROMEGA N3030) following the manufactureŕs protocol. Positive clones were again diluted to single cells to ensure monoclones and 7 positive monoclones were obtained from the screening. For insert validation, gDNA was obtained by using QuickExtract DNA Extraction Solution (Lucigen), where 10000 cells were resuspended in 100 µl PBS, followed by addition of 400 µl QuickExtract and mixing. After 30 minutes of incubation at 65°C, the mix was heat inactivated by a 5 min incubation at 95°C. The DNA in the resulting solution was sheared by pressing the pipet tip to the tube bottom while pipetting up and down and 1 µl was used for subsequent PCR using the primers (fw: CCCTGAAGCTCAGTCTTCTGTCATTTGG; rv: CTACGGGAGGGCTTGCAGAACTG). After PCR cleanup, using the GeneJET PCR purification kit (ThermoFisher K0701), products were sent to sequencing using nested primers (fw: GCACAAACTGCTGCATTCTTACAGACTTC; rv: GGGCAACTATAAAGAGATGTTTTCTGGTTCCTAC). For expression validation, HEK293T wild type cells and cone 4 (positive genomic heterozygous insertion) were used for western blot by running a 12 % SDS PAGE gel, followed by transfer onto a PVDF membrane, using semi-dry blot. The membrane was blocked with 5 % BSA for 1h at RT and WDR5 was detected, using a WDR5 antibody (Santa Cruz Biotechnology, sc-393080) over night at 4 °C with a dilution of 1:200 and subsequent incubation with a HRP-linked secondary anti-mouse antibody (Cell Signaling #7076) for 2h at RT with a 1:5000 dilution. As loading control, Tubulin was detected, using a tubulin antibody (Abcam ab7291) over night at 4°C and HRP linked anti-rabbit secondary antibody (Cell Signaling #7074) for 2h at RT. HRP detection was carried out, using the Clarity Western ECL Blotting Substrate (BioRad) following the manufacturers protocol. For HiBIT detection, the membrane detected, using the Nano-Glo® HiBiT Blotting System (PROMEGA N2410). Here, the LgBIT protein was diluted 1:200 in TBST 5% BSA together with a 1:200 dilution of the substrate solution. Readout for HRP and HiBIT detection was carried out in a ChemiDoc XRS+ (BioRad).

### Western blot for cell line comparison

For expression analysis, HEK293THiBIT-WDR5, HEK293T, MCF7, MV4-11 and THP-1 cells were seeded with 2.5×10^5 cell/ml into 6-well plates and incubated 12 h at 37 °C and 5% CO_2_. After incubation, 5 µM MS67 or MS67N were added to the cells followed by further incubation for 6h at 37 °C and 5 % CO_2_. For cell harvesting medium was aspirated, the cells were washed with cold DPBS (Gibco 14190-094) and 200 µl of cold RIPA buffer (150 mM NaCl, 1 % Triton X-100, 0.1% SDS and 50 mM Tris, pH 8.0) was added to the cells, followed by 20-minute incubation at 4°C. After lysis, the lysate was cleared by centrifugation at 17.000*g* for 30 min at 4 °C. For the SDS PAGE, total protein concentration of the supernatant was measured by BCA assay, using Pierce™ BCA Protein Assay Kit (Thermo Fisher # 23225) with BSA samples for the calibration curve. Here 47.5 µl BCA reagent was mixed with 2.5 µl lysate in duplicates in a clear 348-well plate (Gleiner 781061) and incubated for 20 minutes at 37°C. The readout was carried out in a PHERAstar plate reader (readout at 265 nm). Lysates were diluted to the lowest measured concentration and a NuPAGE grandient gel (4-12 %, Invitrogen NP0321PK2) was run, followed by transfer to a PVDF membrane, using a Trans-Blot Turbo Transfer System (BioRad). The membrane was blocked in TBST 5% BSA for 2h at RT and subsequently treated with TBST, 5% BSA and anti-WDR5 antibody (Santa Cruz Biotechnology, sc-393080) over night at 4 °C with a dilution of 1:200 and subsequent incubation with a HRP-linked secondary anti-mouse antibody (Cell Signaling #7076) for 2h at RT with a 1:5000 dilution. As loading control, Tubulin was detected, using a tubulin antibody (Abcam ab7291) over night at 4°C and HRP linked anti-rabbit secondary antibody (Cell Signaling #7074) for 2h at RT. HRP detection was carried out, using the Clarity Western ECL Blotting Substrate (BioRad) following the manufacturers protocol.

### Western Blot detection of HiBIT tagged WDR5

For visualisation of the protein levels, Western blot was carried out by on-membrane complementation of the Nano Luciferase with subsequent luminescence detection. For this, 2.5×10^5 cell/ml were seeded into 24-well plates and incubated 24 h at 37 °C and 5 % CO2. After incubation, PROTACs were added in different concentrations and the cells were further incubated for the desired time. Compounds were diluted to different concentrations which resulted in addition of 1µl compound in DMSO to 1 ml cells. For cell harvesting medium was aspirated and 50 µl of cold RIPA buffer (150 mM NaCl, 1 % Triton X-100, 0.1% SDS and 50 mM Tris, pH 8.0) was added to the cells, followed by 20-minute incubation at 4°C. For the SDS PAGE, total protein concentration was measured by BCA assay, using Pierce™ BCA Protein Assay Kit (Thermo Fisher # 23225) with BSA samples for the calibration curve. Here 47.5 µl BCA reagent was mixed with 2.5 µl lysate in duplicates in a clear 348-well plate (Gleiner 781061) and incubated for 20 minutes at 37°C. The readout was carried out in a PHERAstar plate reader (readout at 265 nm). Lysates were diluted to the lowest measured concentration and a 12 % SDS PAGE Gel was run, followed by transfer to a PVDF membrane, using a Trans-Blot Turbo Transfer System (BioRad). The membrane was blocked in TBST 5% BSA for 2h at RT and subsequently treated with TBST, 5% BSA and a 1:200 dilutions of LgBIT protein and a 1:200 dilution of the substrate, provided by the Nano-Glo® HiBiT Blotting System (PROMEGA N2410). Readout was carried out in a ChemiDoc XRS+ (BioRad).

### Cell viability assay

PROTAC toxicity was determined by using the CellTiter-Glo® 2.0 Cell Viability Assay (PROMEGA G9241). For this, 40 µl of the CRISPR edited HEK293T cells were seeded with 10000 cells/well into white 384-well plates (Greiner 781207) and the cells were allowed to attach for 24h at 37°C and 5% CO2. After incubation, PROTACs were titrated using an Echo acoustic dispenser (Labcyte) and the plate was further incubated for 24h at 37°C and 5% CO2. For the assay, 40 µl assay reagent was added and incubated for 10 minutes at RT. Filtered luminescence was measured on a PHERAstar plate reader (BMG Labtech) and data was graphed, using GraphPad Prism 9.

### In-cell ELISA

For immunofluorescence staining of WDR5 and Tubulin, HEK293T and HEK293T^HiBIT-WDR5^ cells were seeded into black 384-well clear bottom plates (Greiner 781866) with 3000 cells per well. The cells containing the LgBIT protein, 2,5 µl transfection reagent, prepared following the FuGENE HD (PROMEGA) protocol was added to 50 µl of seeded cells. After seeding, the cells were incubated overnight at 37 °C and 5% CO2 to allow the cells to attach. After attachment, the cells were washed with PBS and fixed with 100 µl 4% paraformaldehyde in PBS for 30 min followed by a wash step with PBS and cell permeabilization in 0.1% Triton X-100 in PBS for 30 min. For blocking, 5% BSA in TBST was used and incubated for 2h at RT. As primary antibodies, anti-Tubulin (Abcam, ab6046) and anti WDR5 (Santa Cruz Biotechnology, sc-393080) were used in 1:1000 and 1:100 dilutions, respectively and incubated overnight. Wells were washed with TBST and secondary antibodies (anti-Rb: anti-Ms: Abcam, ab150113) were used in 1:1000 dilution and incubated for 2h at RT while shaking at 400 rpm in the dark. Finally, Hoechst33342 (Thermo Fisher; H3570) staining was carried out by using 1 µM stain and incubate additional 15 minutes at RT while shaking. Imaging was carried out in a Cellcyte X (Cytena) by detection of bright field, red (Tubulin), blue (Hoechst33342) and green (WDR5) channel.

### NanoBRET assay

Full-length VHL and WDR5 were obtained as plasmids cloned in frame with a terminal NanoLuc-fusion (VHL: Promega, ND2700). The plasmid was transfected into HEK293T cells using FuGENE HD (Promega, E2312) and proteins were allowed to express for 20h. Serially diluted inhibitor and NanoBRET VHL Tracer (Promega) or WDR5 tracer (19c Dölle *et al.*) at a concentration previously determined (1 µM) were pipetted into white 384-well plates (Greiner 781207) using an Echo acoustic dispenser (Labcyte). The corresponding protein-transfected cells were added and reseeded at a density of 2 x 10^5^ cells/ml after trypsinization and resuspending in Opti-MEM without phenol red (Life Technologies). The system was allowed to equilibrate for 2 hours at 37°C/5% CO2 prior to BRET measurements. To measure BRET, NanoBRET NanoGlo Substrate + Extracellular NanoLuc Inhibitor (Promega, N2540) were added as per the manufacturer’s protocol, and filtered luminescence was measured on a PHERAstar FSX plate reader (BMG Labtech) equipped with a luminescence filter pair (450 nm BP filter (donor) and 610 nm LP filter (acceptor)). Competitive displacement data were then graphed using GraphPad Prism 9 software using a normalized 3-parameter curve fit with the following equation: Y=100/(1+10^(X-LogIC_50_)).

### HiBIT-based intact cell CETSA

For intact cell melting point determination, HEK293T^HiBIT-WDR5^ were used in a concentration of 200000 cells/ml, resuspended in OptiMem medium. The cells were incubated with 10 µM of the corresponding compound for 1h at 37°C and 5% CO2. For lysed cell assay, additional 50 ng/µl digitonin was added with the compounds followed by incubation at RT for 30 min. Incubated cells were transferred into 96-well PCR plates (STARLAB; I1402-9800) and incubated for 3 min in a gradient PCR cycler (Biometra), followed by 3 min cool down at 25 °C. 10 µl of the incubated cells were transferred into white 384-well LDV plates (Greiner 784075) and 10 µl HiBIT lytic detection reagent was added following the manufactureŕs protocol. Readout was carried out in a PheraStar FSX plate reader (BMG Labtech) using the LUM plus optical module while the plate was measured for 10 minutes every minute. Data was graphed, using GraphPad Prism 9 and cellular melting points were determined by the melting curves IC50, estimated through a four parameter fit with the equation Y=Bottom + (Top-Bottom)/(1+(IC_50_/X)^HillSlope).

### Cellular ternary complex formation measurement

For ternary complex measurements, WDR5 and VHL were cloned in frame with C and N-terminal NanoLuc or HaloTag, respectively to generate every combination. All combinations were tested for the identification of the pair which provides the best signal which were found to be VHL-NanoLuc and WDR5-HaloTag. These plasmids were subsequently used for all measurements. For the transfections, HEK293T cells were diluted in OptiMEM without phenol red (Life Technologies) to 4 x 10^5^ cells/ml and 38 µl were pipetted into white 384-well plates (Greiner 781207). 1 ml FuGENE HD (Promega, E2312) transfection mix containing 30 µl FuGENE HD and 8 µl of each plasmid was prepared per plate, incubated 20 min at RT and 2 µl were pipetted into each well leading to the total assay volume of 40 µl. After incubation at 37 °C/5 % CO_2_ for 20 h 40 nl HaloTag® NanoBRET™ 618 Ligand (PROMEGA) was added to the cells using an Echo acoustic dispenser (Labcyte) and the cells were incubated additional 20h at 37 °C/5 % CO_2_. 2 h prior BRET measurement, the compounds were titrated to the cells using an Echo acoustic dispenser and the cells were incubated further at 37 °C/5 % CO_2_ to allow complex formation. To measure BRET, NanoBRET NanoGlo Substrate + Extracellular NanoLuc Inhibitor (Promega, N2540) was added as per the manufacturer’s protocol, and filtered luminescence was measured on a PHERAstar FSX plate reader (BMG Labtech) equipped with a luminescence filter pair (450 nm BP filter (donor) and 610 nm LP filter (acceptor)). Stimulation data were then graphed using GraphPad Prism 9 software using a normalized 3-parameter curve fit with the following equation: Y=Bottom + (Top-Bottom)/(1+10^((LogEC50-X))).

### Measurement of cellular ternary complex kinetics

Kinetics measurement were carried out, using the C-terminally Nano Luciferase tagged VHL and C-terminally HaloTagged WDR5. Transfection and incubation were carried out as described above. After the addition of the HaloTag® NanoBRET™ 618 Ligand and 20h incubation, NanoBRET NanoGlo Substrate + Extracellular NanoLuc Inhibitor (Promega, N2540) was added as per the manufacturer’s protocol. The kinetics measurement was started with PROTAC addition and filtered luminescence was measured on a PHERAstar FSX plate reader (BMG Labtech) equipped with a luminescence filter pair (450 nm BP filter (donor) and 610 nm LP filter (acceptor)). PROTACs were used in a first round in 5 µM to estimate IC_50_ values and measurements were carried out with 10-fold PROTAC access (SI Table 3), resulting in comparable IC_50_ values. After 1h measurement, the wells were washed with PBS and fresh OptiMEM containing substrate and NanoLuc inhibitor was added for dissociation measurement and the measurement was carried out for additional 3h. Ratio of BRET and luminescence was calculated and data was graphed using GraphPad Prism 9. Half-time was calculated, using a one phase exponential decay using following equation (Y=(Y0-NS)*exp(-K*X) + NS).

### Cellular degradation kinetics

For cellular degradation kinetics, HEK293T^HiBIT-WDR5^ cells were transfected in a T75 flask, using FuGENE 4K (PROMEGA) following the manufactureŕs protocol with the LgBIT. After allowing the cells to express the LgBIT protein for 20h at 37 °C and 5% CO2, the cells were harvested and the medium was exchanged to CO2 independent medium (Life Sciences) containing Endurazine substrate (PROMEGA) according to the manufacturers protocol. The cells were either transferred into a 96-well (100 µl) or 384-well (40 µl) plate and incubated for 2.5h prior readout for substrate activation. After incubation, compounds were added and the plate was sealed with BreathEasy plate seal (Sigma Aldrich Z380059) and continuously measured every 5 min for 24/60h in a PHERAstar FSX plate reader (BMG Labtech) using the LUM plus module. Untreated baseline measurements were subtracted from the measurements to normalize the data and graphed, using GraphPad Prism 9.

### Protein Purification

WDR5 and the complex of VHL-EloB-EloC were expressed and purified as previously described.^21,30^

For WDR5, the pET28a-LIC plasmid containing WDR5_23-334_ along with a N-terminal 6×His Tag and thrombin cutting site was used and a kind gift of M. Vedadi (SGC Toronto). WDR5 was overexpressed in Escherichia coli BL21 (DE3) using TB media supplemented with 50 µg/ml kanamycin. Protein expression was induced by addition of 0.5 mM IPTG at an OD_600_ of approx. 3. Cells were grown overnight at 18 °C. Next morning, the cells were harvested for 10 minutes at 6000 rpm (Thermo Scientific™ Sorvall™ LYNX™) Superspeed and resuspended in Lysis buffer (50 mM HEPES buffer, pH 7.5, 500 mM NaCl, 20 mM imidazole, 0.5 mM TCEP, and 5% glycerol). For purification, the cells were lysed by sonication. After centrifugation at 20.000 rpm for 30 minutes, the supernatant was loaded onto a Nickel−Sepharose column equilibrated with 30 mL of lysis buffer. The column was washed with 100 mL of lysis buffer. WDR5 was eluted by an imidazole step gradient (50, 100, 200, 300 mM). Fractions containing WDR5 were pooled together, concentrated, and loaded onto a Superdex 200 16/ 60 HiLoad (GE Healthcare) gel filtration column equilibrated with final buffer (25 mM HEPES pH 7.5, 300 mM NaCl, and 0.5 mM TCEP). The protein was concentrated to approx. 25 mg/mL, frozen in liquid nitrogen and stored at minus 80°C.

For VHL, the pHAT4 plasmid containing VHL_54-213_ along with a N-terminal 6×His Tag and TEV cutting site and the pDUET-1 plasmid encoding for both EloB_1-120_ and EloC_17-112_ were used and a kind gift from A. Ciulli (University of Dundee). The VHL-ElonginB-ElonginC complex was co-expressed in Escherichia coli BL21 (DE3) using TB media supplemented with 100 μg/ml ampicillin and 50 μg/ml streptomycin. Protein expression was induced by addition of 0.5 mM IPTG at an OD_600_ of approx. 3. Cells were grown overnight at 18 °C. Next morning, the cells were harvested for 10 minutes at 6000 rpm and resuspended in Lysis buffer (50 mM HEPES buffer, pH 7.5, 500 mM NaCl, 20 mM imidazole, 0.5 mM TCEP, and 5% glycerol). For purification, the cells were lysed by sonication. After centrifugation at20.000 rpm for 30 minutes, the supernatant was loaded onto a Nickel−Sepharose column equilibrated with 30 mL of lysis buffer. The column was washed with 100 mL of lysis buffer. The protein complex was eluted by an imidazole step gradient (50, 100, 200, 300 mM). To remove the expression tag on VHL, the fractions containing the complex were combined, supplemented with TEV protease (ratio approx. 20:1) and dialysed against final buffer (25 mM HEPES pH 7.5, 200 mM NaCl, and 0.5 mM TCEP) overnight at 4°C. Next day a reverse Nickel−Sepharose column equilibrated with final buffer was used to remove uncleaved protein and tag. The complex which was mainly in the flow through fraction was concentrated to approx. 5 mL and loaded onto a Superdex 200 16/ 60 HiLoad gel filtration column equilibrated with a final buffer. The complex was concentrated to approx. 15 mg/mL, frozen in liquid nitrogen and stored at minus 80°C.

### Crystallization of ternary complex

WDR5, VHL-EloB-EloC, and the desired PROTAC (10 mM in DMSO) were mixed in a 1:1:1 ratio and incubated for 30 min on ice. The final concentration of each component was approx. 100 µM. Crystals were obtained using the sitting drop vapour diffusion method at 20°C (SWISS CI 96 well 3 lens plate). The reservoir condition contained 0.1 M HEPES (pH 6.8-7.5), 0.4-1.2 M sodium thiocyanate, and 18-20% PEG 3350.

The crystallization of a ternary complex with JW48 was not successful. Obtained crystals were solved to be a binary complex with VHL-EloB-EloC. Crystals grow in 19% PEG 3350, 0.1 M HEPES pH7.

### Data collection, structure solution and refinement

Diffraction data were collected at beamline X06SA or X06DA (Villigen, CH) at a wavelength of 1.0 Å at 100 K. The reservoir condition supplemented with 25% ethylene glycol was used as cryoprotectant. Data were processed using XDS^31^ and scaled with aimless.^32^ The PDB structures with the accession code 2GNQ^33^ (for WDR5) and 1VCB^34^ (for VHL-EloB-EloC) were used as initial search models using the program PHASER.^35^ The final model was built manually using Coot^36^ and stepwise refined with REFMAC5.^37^ Data collection and refinement statistics are summarized in SI Table 6.

### In vitro NanoBRET assay

Purified HiBIT-tagged full length WDR5 (5 nM) was mixed with a 1:100 dilution of LgBIT protein (Promega) in DPBS (Gibco 14190-094). The mixture was used for assays in 10 µl in white 384-LDV plates (Greiner 784075) where the tracer KD was previously determined by a tracer titration using the WDR5 tracer (19c Dölle *et al.*) and found to be 30 nM. For readout, 5 µl of NanoBRET NanoGlo Substrate in DPBS (Gibco 14190-094) was added to the wells followed by readout in a PHERAstar FSX plate reader (BMG Labtech) equipped with a luminescence filter pair (450 nm BP filter (donor) and 610 nm LP filter (acceptor)). For dose-response measurements, compounds were titrated to 5 nM HiBIT-WDR5, LgBIT and 30 nM of WDR5 tracer followed by addition of 5 µl NanoBRET NanoGlo Substrate in DPBS (Gibco 14190-094). Filtered luminescence was measured on a PHERAstar FSX plate reader (BMG Labtech) equipped with a luminescence filter pair (450 nm BP filter (donor) and 610 nm LP filter (acceptor)). Competitive displacement data were then graphed using GraphPad Prism 9 software using a normalized 3-parameter curve fit with the following equation: Y=100/(1+10^(X-LogIC50)).

### Differential scanning fluorimetry

For melting temperature determination, purified proteins were buffered in DPBS (Gibco 14190-094) and were assayed in a 384-well plate (Thermo, #BC3384) with a final protein concentration of 5 μM in 10 μL final assay volume. As a fluorescent probe, SYPRO-Orange (Molecular Probes) was used in a 1:1000 dilution. Filters for excitation and emission were set to 465 and 590 nm, respectively. The temperature was increased from 25°C with 3°C/min to a final temperature of 99°C, while scanning, using the QuantStudio5 (Applied Biosystems). Data was analyzed using Boltzmannequation in the Protein Thermal Shift software (Applied Biosystems). Samples were measured in technical quadruplikates.

### Measurement of *in vitro* ternary complexes through mass photometry

Purified WDR5 and VHL-EloB-EloC was used for mass photometry experiments, together with the PROTACs with available crystal structures (AD122, AD142, AD157 and MS67) together with JW48. For all measurements, a REFEYN Two^MP^ was used. High precision microscope cover glasses (Thorlabs CG15KH) were cleaned and prepared with Grace Bio-Labs CultureWell gaskets (Sigma Aldrich GBL103350) to allow the measurement of 6 drops. 15 µl dilution buffer (25 mM HEPES pH 7.5, 200 mM NaCl, and 0.5 mM TCEP, 5% Glycerol) were used for the autofocus followed by in-drop dilution of the purified proteins. For this, the proteins were diluted in dilution buffer to a concentration of 100 nM and again diluted in the drop 4-fold by adding 5 µL of dilution buffer leading to a final concentration of 25 nM. For data collection, videos were recorded for 1 min. For the calibration curve, Carbonic Anhydrase (29 kDa), albumin (66 kDa) and beta-amylase (56, 112 and 224) were used and obtained as a kind gift from Dr Mohit Misra. For ternary complex formation assays, either 10 µM PROTAC were used for absolute complex validation or a dose response (10 µM, 3 µM, 920 nM and 280 nM) was measured for MS67. After mixing VHL-EloB-EloC, WDR5 and the PROTACs, the samples were incubated on ice for 15 minutes to allow the complexes to form. Results were analyzed and graphed in Refeyn DiscoverMP using the measured calibration curve.

## References

1. Nemec, V., Schwalm, M.P., Muller, S., and Knapp, S. (2022). PROTAC degraders as chemical probes for studying target biology and target validation. Chem Soc Rev 51, 7971–7993. 10.1039/d2cs00478j.

2. Li, K., and Crews, C.M. (2022). PROTACs: past, present and future. Chem Soc Rev 51, 5214–5236. 10.1039/d2cs00193d.

3. Schwalm, M.P., and Knapp, S. (2022). BET bromodomain inhibitors. Curr Opin Chem Biol 68, 102148. 10.1016/j.cbpa.2022.102148.

4. Troup, R.I., Fallan, C., and Baud, M.G.J. (2020). Current strategies for the design of PROTAC linkers: a critical review. Explor Target Antitumor Ther 1, 273–312. 10.37349/etat.2020.00018.

5. Roy, M.J., Winkler, S., Hughes, S.J., Whitworth, C., Galant, M., Farnaby, W., Rumpel, K., and Ciulli, A. (2019). SPR-Measured Dissociation Kinetics of PROTAC Ternary Complexes Influence Target Degradation Rate. ACS Chem Biol 14, 361–368. 10.1021/acschembio.9b00092.

6. Smith, B.E., Wang, S.L., Jaime-Figueroa, S., Harbin, A., Wang, J., Hamman, B.D., and Crews, C.M. (2019). Differential PROTAC substrate specificity dictated by orientation of recruited E3 ligase. Nat Commun 10, 131. 10.1038/s41467-018-08027-7.

7. Buckley, D.L., Raina, K., Darricarrere, N., Hines, J., Gustafson, J.L., Smith, I.E., Miah, A.H., Harling, J.D., and Crews, C.M. (2015). HaloPROTACS: Use of Small Molecule PROTACs to Induce Degradation of HaloTag Fusion Proteins. ACS Chem Biol 10, 1831–1837. 10.1021/acschembio.5b00442.

8. Schwinn, M.K., Machleidt, T., Zimmerman, K., Eggers, C.T., Dixon, A.S., Hurst, R., Hall, M.P., Encell, L.P., Binkowski, B.F., and Wood, K.V. (2018). CRISPR-Mediated Tagging of Endogenous Proteins with a Luminescent Peptide. ACS Chem Biol 13, 467–474. 10.1021/acschembio.7b00549.

9. Riching, K.M., Mahan, S., Corona, C.R., McDougall, M., Vasta, J.D., Robers, M.B., Urh, M., and Daniels, D.L. (2018). Quantitative Live-Cell Kinetic Degradation and Mechanistic Profiling of PROTAC Mode of Action. ACS Chem Biol 13, 2758–2770. 10.1021/acschembio.8b00692.

10. Schapira, M., Tyers, M., Torrent, M., and Arrowsmith, C.H. (2017). WD40 repeat domain proteins: a novel target class? Nat Rev Drug Discov 16, 773–786. 10.1038/nrd.2017.179.

11. Chen, X., Xu, J., Wang, X., Long, G., You, Q., and Guo, X. (2021). Targeting WD Repeat-Containing Protein 5 (WDR5): A Medicinal Chemistry Perspective. J Med Chem 64, 10537–10556. 10.1021/acs.jmedchem.1c00037.

12. Su, J., Wang, F., Cai, Y., and Jin, J. (2016). The Functional Analysis of Histone Acetyltransferase MOF in Tumorigenesis. Int J Mol Sci 17. 10.3390/ijms17010099.

13. Bode, D., Yu, L., Tate, P., Pardo, M., and Choudhary, J. (2016). Characterization of Two Distinct Nucleosome Remodeling and Deacetylase (NuRD) Complex Assemblies in Embryonic Stem Cells. Mol Cell Proteomics 15, 878–891. 10.1074/mcp.M115.053207.

14. Guarnaccia, A.D., and Tansey, W.P. (2018). Moonlighting with WDR5: A Cellular Multitasker. J Clin Med 7. 10.3390/jcm7020021.

15. Li, Y., Han, J., Zhang, Y., Cao, F., Liu, Z., Li, S., Wu, J., Hu, C., Wang, Y., Shuai, J., et al. (2016). Structural basis for activity regulation of MLL family methyltransferases. Nature 530, 447–452. 10.1038/nature16952.

16. Senisterra, G., Wu, H., Allali-Hassani, A., Wasney, G.A., Barsyte-Lovejoy, D., Dombrovski, L., Dong, A., Nguyen, K.T., Smil, D., Bolshan, Y., et al. (2013). Small-molecule inhibition of MLL activity by disruption of its interaction with WDR5. Biochem J 449, 151–159. 10.1042/BJ20121280.

17. Karatas, H., Li, Y., Liu, L., Ji, J., Lee, S., Chen, Y., Yang, J., Huang, L., Bernard, D., Xu, J., et al. (2017). Discovery of a Highly Potent, Cell-Permeable Macrocyclic Peptidomimetic (MM-589) Targeting the WD Repeat Domain 5 Protein (WDR5)-Mixed Lineage Leukemia (MLL) Protein-Protein Interaction. J Med Chem 60, 4818-4839. 10.1021/acs.jmedchem.6b01796.

18. Macdonald, J.D., Chacon Simon, S., Han, C., Wang, F., Shaw, J.G., Howes, J.E., Sai, J., Yuh, J.P., Camper, D., Alicie, B.M., et al. (2019). Discovery and Optimization of Salicylic Acid-Derived Sulfonamide Inhibitors of the WD Repeat-Containing Protein 5-MYC Protein-Protein Interaction. J Med Chem 62, 11232–11259. 10.1021/acs.jmedchem.9b01411.

19. Cao, F., Townsend, E.C., Karatas, H., Xu, J., Li, L., Lee, S., Liu, L., Chen, Y., Ouillette, P., Zhu, J., et al. (2014). Targeting MLL1 H3K4 methyltransferase activity in mixed-lineage leukemia. Mol Cell 53, 247–261. 10.1016/j.molcel.2013.12.001.

20. Yu, X., Li, D., Kottur, J., Shen, Y., Kim, H.S., Park, K.S., Tsai, Y.H., Gong, W., Wang, J., Suzuki, K., et al. (2021). A selective WDR5 degrader inhibits acute myeloid leukemia in patient-derived mouse models. Sci Transl Med 13, eabj1578. 10.1126/scitranslmed.abj1578.

21. Dolle, A., Adhikari, B., Kramer, A., Weckesser, J., Berner, N., Berger, L.M., Diebold, M., Szewczyk, M.M., Barsyte-Lovejoy, D., Arrowsmith, C.H., et al. (2021). Design, Synthesis, and Evaluation of WD-Repeat-Containing Protein 5 (WDR5) Degraders. J Med Chem 64, 10682-10710. 10.1021/acs.jmedchem.1c00146.

22. Mortison, J.D., Cornella-Taracido, I., Venkatchalam, G., Partridge, A.W., Siriwardana, N., and Bushell, S.M. (2021). Rapid Evaluation of Small Molecule Cellular Target Engagement with a Luminescent Thermal Shift Assay. ACS Med Chem Lett 12, 1288–1294. 10.1021/acsmedchemlett.1c00276.

23. Roy, M.J., Winkler, S., Hughes, S.J., Whitworth, C., Galant, M., Farnaby, W., Rumpel, K., and Ciulli, A. (2019). SPR-measured dissociation kinetics of PROTAC ternary complexes influence target degradation rate. ACS chemical biology 14, 361–368.

24. Cao, C., He, M., Wang, L., He, Y., and Rao, Y. (2022). Chemistries of bifunctional PROTAC degraders. Chem Soc Rev 51, 7066–7114. 10.1039/d2cs00220e.

25. Donovan, K.A., Ferguson, F.M., Bushman, J.W., Eleuteri, N.A., Bhunia, D., Ryu, S., Tan, L., Shi, K., Yue, H., Liu, X., et al. (2020). Mapping the Degradable Kinome Provides a Resource for Expedited Degrader Development. Cell 183, 1714–1731 e1710. 10.1016/j.cell.2020.10.038.

26. Douglass, E.F., Jr., Miller, C.J., Sparer, G., Shapiro, H., and Spiegel, D.A. (2013). A comprehensive mathematical model for three-body binding equilibria. J Am Chem Soc 135, 6092–6099. 10.1021/ja311795d.

27. Krissinel, E., and Henrick, K. (2007). Inference of macromolecular assemblies from crystalline state. J Mol Biol 372, 774–797. 10.1016/j.jmb.2007.05.022.

28. Yu, X., Li, D., Kottur, J., Shen, Y., Kim, H.S., Park, K.S., Tsai, Y.H., Gong, W., Wang, J., Suzuki, K., et al. (2021). A selective WDR5 degrader inhibits acute myeloid leukemia in patient-derived mouse models. Science translational medicine 13, eabj1578. 10.1126/scitranslmed.abj1578.

29. Wang, R., Ascanelli, C., Abdelbaki, A., Fung, A., Rasmusson, T., Michaelides, I., Roberts, K., and Lindon, C. (2021). Selective targeting of non-centrosomal AURKA functions through use of a targeted protein degradation tool. Commun Biol 4, 640. 10.1038/s42003-021-02158-2.

30. Buckley, D.L., Van Molle, I., Gareiss, P.C., Tae, H.S., Michel, J., Noblin, D.J., Jorgensen, W.L., Ciulli, A., and Crews, C.M. (2012). Targeting the von Hippel-Lindau E3 ubiquitin ligase using small molecules to disrupt the VHL/HIF-1alpha interaction. J Am Chem Soc 134, 4465–4468. 10.1021/ja209924v.

31. Kabsch, W. (2010). Xds. Acta Crystallogr D Biol Crystallogr 66, 125–132. 10.1107/S0907444909047337.

32. Evans, P.R., and Murshudov, G.N. (2013). How good are my data and what is the resolution? Acta Crystallogr D Biol Crystallogr 69, 1204–1214. 10.1107/S0907444913000061.

33. Schuetz, A., Allali-Hassani, A., Martin, F., Loppnau, P., Vedadi, M., Bochkarev, A., Plotnikov, A.N., Arrowsmith, C.H., and Min, J. (2006). Structural basis for molecular recognition and presentation of histone H3 by WDR5. EMBO J 25, 4245–4252. 10.1038/sj.emboj.7601316.

34. Stebbins, C.E., Kaelin, W.G., Jr., and Pavletich, N.P. (1999). Structure of the VHL-ElonginC-ElonginB complex: implications for VHL tumor suppressor function. Science 284, 455–461. 10.1126/science.284.5413.455.

35. McCoy, A.J., Grosse-Kunstleve, R.W., Adams, P.D., Winn, M.D., Storoni, L.C., and Read, R.J. (2007). Phaser crystallographic software. J Appl Crystallogr 40, 658–674. 10.1107/S0021889807021206.

36. Emsley, P., and Cowtan, K. (2004). Coot: model-building tools for molecular graphics. Acta Crystallogr D Biol Crystallogr 60, 2126–2132. 10.1107/S0907444904019158.

37. Vagin, A.A., Steiner, R.A., Lebedev, A.A., Potterton, L., McNicholas, S., Long, F., and Murshudov, G.N. (2004). REFMAC5 dictionary: organization of prior chemical knowledge and guidelines for its use. Acta Crystallogr D Biol Crystallogr 60, 2184–2195. 10.1107/S0907444904023510.

