## Supplementary File for "Tracking the PROTAC degradation pathway in living cells highlights the importance of ternary complex measurement for PROTAC optimization"

### WDR5 HiBIT insertion validation through sanger sequencing:

For N-terminal HiBIT fusion, the canonical WDR5 protein (UniProt: P61964) was used as reference protein sequence. Here, all isoforms start with Exon 2, encoding MATEEKKPETEAARAQPTPSSSATQSK, while Exon 1 is coding for AAWRPPELPPCRAESALPPRRRPTRRPERPAPPPRPG, a sequence not found in the isoforms reportet on UniProt. The guide RNA was designed, based on the CRISPR tool of www.benchling.com.(GCTGGCCGCACAGGAGACAA).

HiBIT-tagged WDR5 cell line through CRISPR/Cas9 was generated as described in the Material and Methods section. After isolation of 7 monoclones from the cell pool, western blot analysis was run to identify clones which were positive for LgBIT complementation in western blot (SI Figure 1).

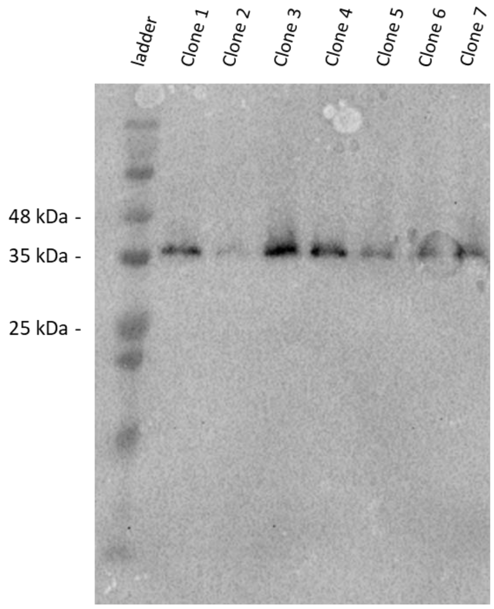


**SI Figure 1: Membrane detection after LgBIT treatment and addition of the luciferase substrate.**

Clones were used for gDNA isolation and nested PCR was used for subsequent sanger sequencing. Generally, the reads were moderate and have shown mixed signals for the locations, indicating only heterozygous clones. From all clones, clone 4 has shown the highest signals for the correct bases (SI Figure 2).

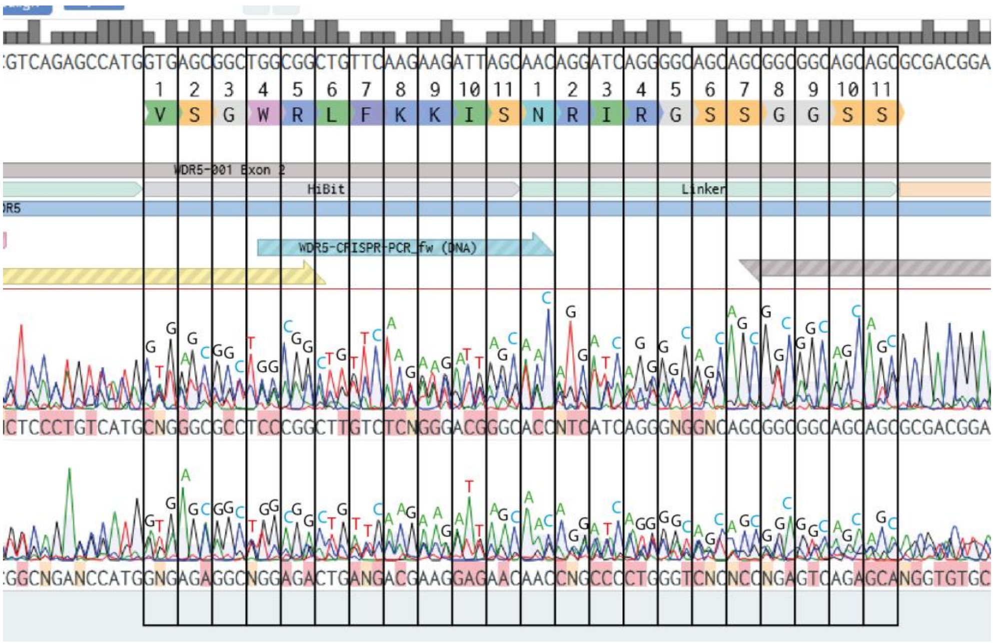


**SI Figure 2: Sequencing alignment of clone 4 towards the insertion site**

Despite the mixed signals (due to heterozygous gDNA), all bases of the insertion are clearly present, proving the insertion in addition to the western blot.

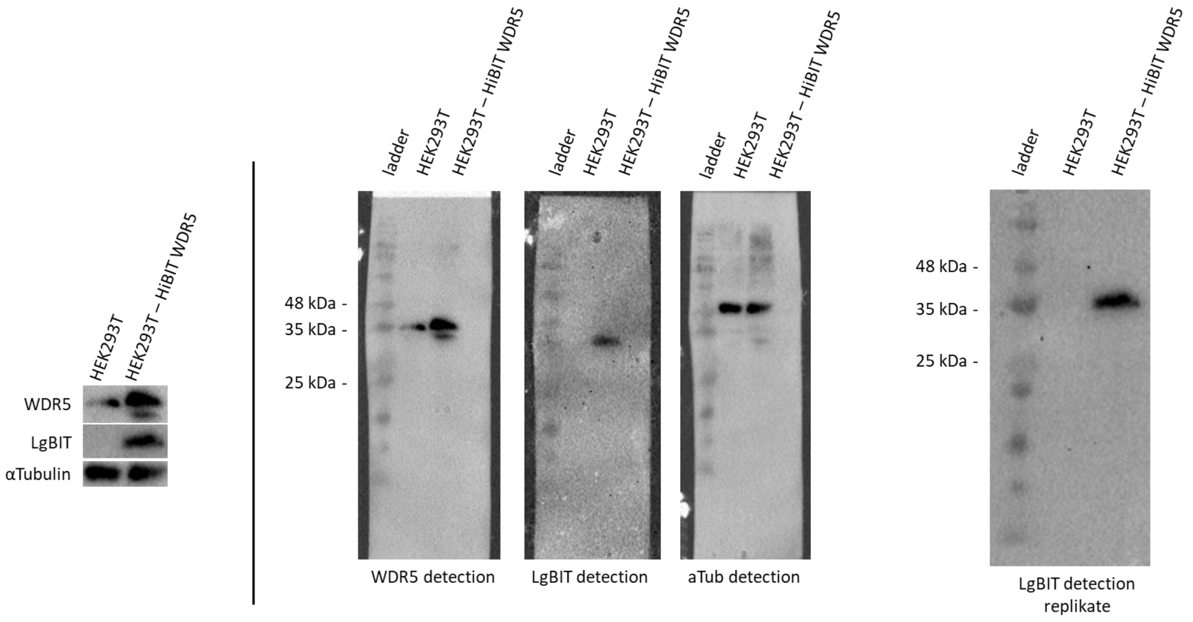


**SI Figure 3: Western blot validation of clone 4**

Western blot on clone 4 shows only a single signal for HiBIT detection, proving only a single insertion at the right size, fitting to the corresponding WDR5 size. A second WDR5 band is visible using WDR5 antibodies which is not showing a signal in LgBIT detection, therefore lacking the HiBIT insertion.

All experiments prove that there was a heterozygous insertion of the HiBIT to the N-terminus of WDR5 in the used HEK293T cells. However, an artefact of WDR5 was created (smaller size) which anyhow should not influence overall protein level measurements.

**Protein expression comparison:**

Relative expression levels of WDR5-PROTAC relevant proteins in leukemia cell lines in comparison to HEK293T cells.

**SI-Table 1: Relative protein levels present in HEK293T cells in comparison to different cell lines in %**

| | | WDR5 | VHL | EloB | EloC | Cullin2 | Rbx1 | CDC34 | UBCH5A | MYC | PAXBP1 | MLL1 | MLL2 |
|---|---|---|---|---|---|---|---|---|---|---|---|---|---|
| HL-60 | Acute promyelocytic leukemia | 50,66666667 | 71,2473573 | 214,156735 | 117,524917 | 125,524476 | 117,801753 | 374,358974 | 138,562092 | 59,2198582 | 54,1666667 | 35,0877193 | 78,1609195 |
| THP-1 | Acute monocytic leukemia | 58,23754789 | 57,0219966 | 117,124235 | 110,633307 | 110,802469 | 175,577889 | 254,355401 | 207,843137 | 96,9802555 | 97,3262032 | 109,89011 | 63,8497653 |
| K-562 | Chronic Myelogenous Leukemia | 60,55776892 | 69,7722567 | 229,489164 | 85,1383875 | 59,338843 | 241,966759 | 121,363259 | 212 | 30,3746817 | 53,0612245 | 30,4878049 | 60,4444444 |
| MOLT-4 | Acute lymphoblastic leukemia | 58,91472868 | 74,0659341 | 128,997172 | 45,3962143 | 75,2620545 | 193,68071 | 273,921201 | 128,484848 | 42,3427992 | 78,7878788 | 54,9450549 | 85,5345912 |
| HEL | Erythroleukemia | 114,861461 | 63,9468691 | 314,588859 | 76,1981691 | 69,3050193 | 289,238411 | 525,179856 | 432,653061 | 45,0499056 | 61,4864865 | 21,8818381 | 45,7912458 |
| HMC-1 | Mast cell leukaemia | 113,4328358 | 81,598063 | 206,908583 | 105,913174 | 110,122699 | 233,55615 | 450,617284 | 179,661017 | 321,153846 | 106,432749 | 128,205128 | 52,1072797 |
| MCF7 | metastatic adenocarcinoma | 39,27648579 | 96,2857143 | 48,8951187 | 90,7633098 | 104,360465 | 127,798098 | 180,916976 | 347,540984 | 109,94075 | 144,444444 | 78,7401575 | 39,8826979 |

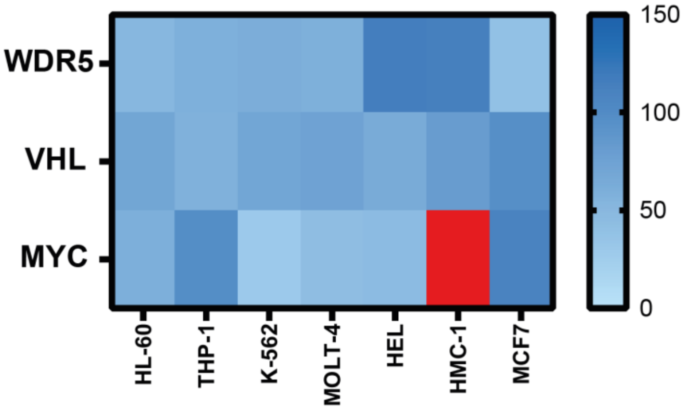

**SI Figure 4: Relative protein expression levels of WDR5, VHL and MYC in leukemia cell lines in comparison to HEK293T cells according to Human Protein Atlas (PMID: 25613900**). Red = > 150

**In-cell ELISA:**

In addition to the unmodified HEK293T and HEK293T$^{HiBIT\text{-}WDR5}$ cells, LgBIT transfected cells were measured in the in-cell ELISA as well, to show no change in localization.

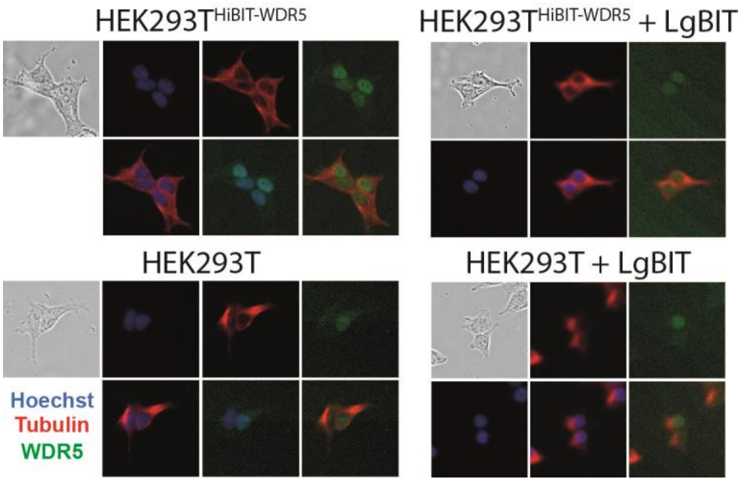


**SI-Figure 5: In-cell ELISA results for unmodified cell lines and cells lines ectopically expressing the LgBIT**. Hoechst staining (blue) = nucleus, Tubulin staining (red) = cytosol, WDR5 staining (green) = WDR5 cellular localization

**Endogenous NanoBRET:**

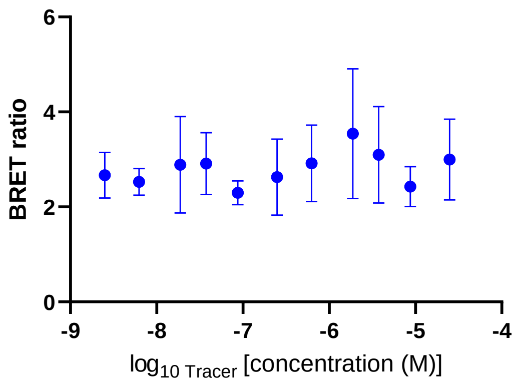


**SI Figure 6: Tracertitration of the WDR5 tracer in HEK293T$^{HiBIT\text{-}WDR5}$ cells expressing the LgBIT.** Average luciferase signal was measured to be 115990 proving successful transfection and complementation of the LgBIT, excluding this as failure reason for the assay. The graph shows error bars based on the standard deviation of two technical duplikates. n=2

**Cell viability determination:**

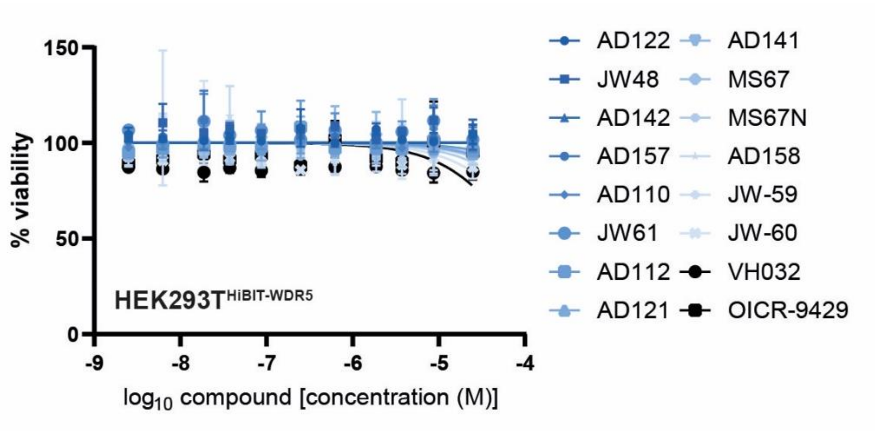


**SI Figure 7: Cell viability after 24h treatment with the different compounds in dose response.** The graph shows error bars based on the standard deviation of biological duplicates. n=2

**Intact Cell CETSA:**

For the establishment of the cellular thermal shift assay for endogenous HiBIT-tagged WDR5, different assay parameters were tested, resulting in the best fitting parameters, described in the material and methods part. The incubation time of the compounds was expected to be a crucial step to allow complete occupation of the binding site. For this, the establishment assay using OICR-9429 and DMSO were tested and incubated either one or two hours with subsequent readout.

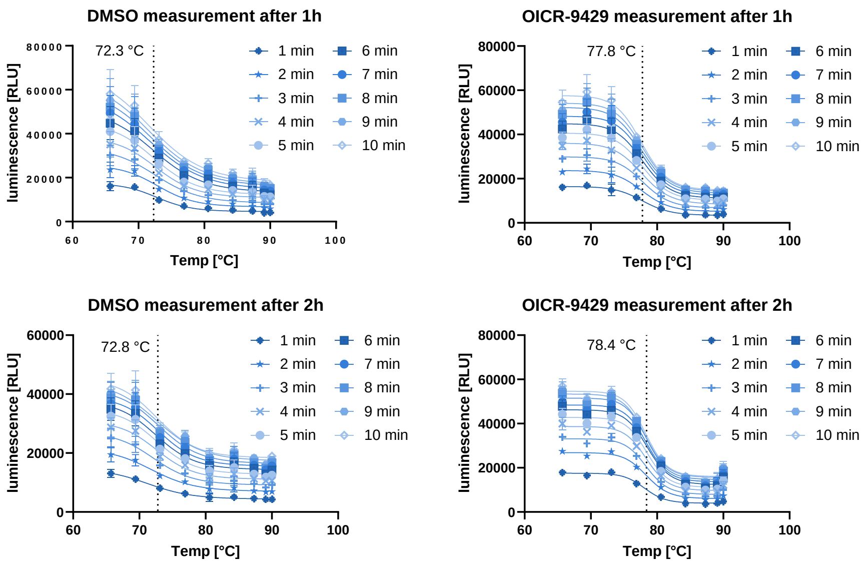


**SI Figure 8: HiBIT-based CETSA data collected after 1 and 2h incubation with either DMSO or OICR-9429.** Data was collected for 10 minutes after addition of the lysis buffer every minute to track the signal during lysis. Calculated melting temperatures correlated during increasing time. Graphs show error bars based on the standard deviation of two technical duplikates. n=2

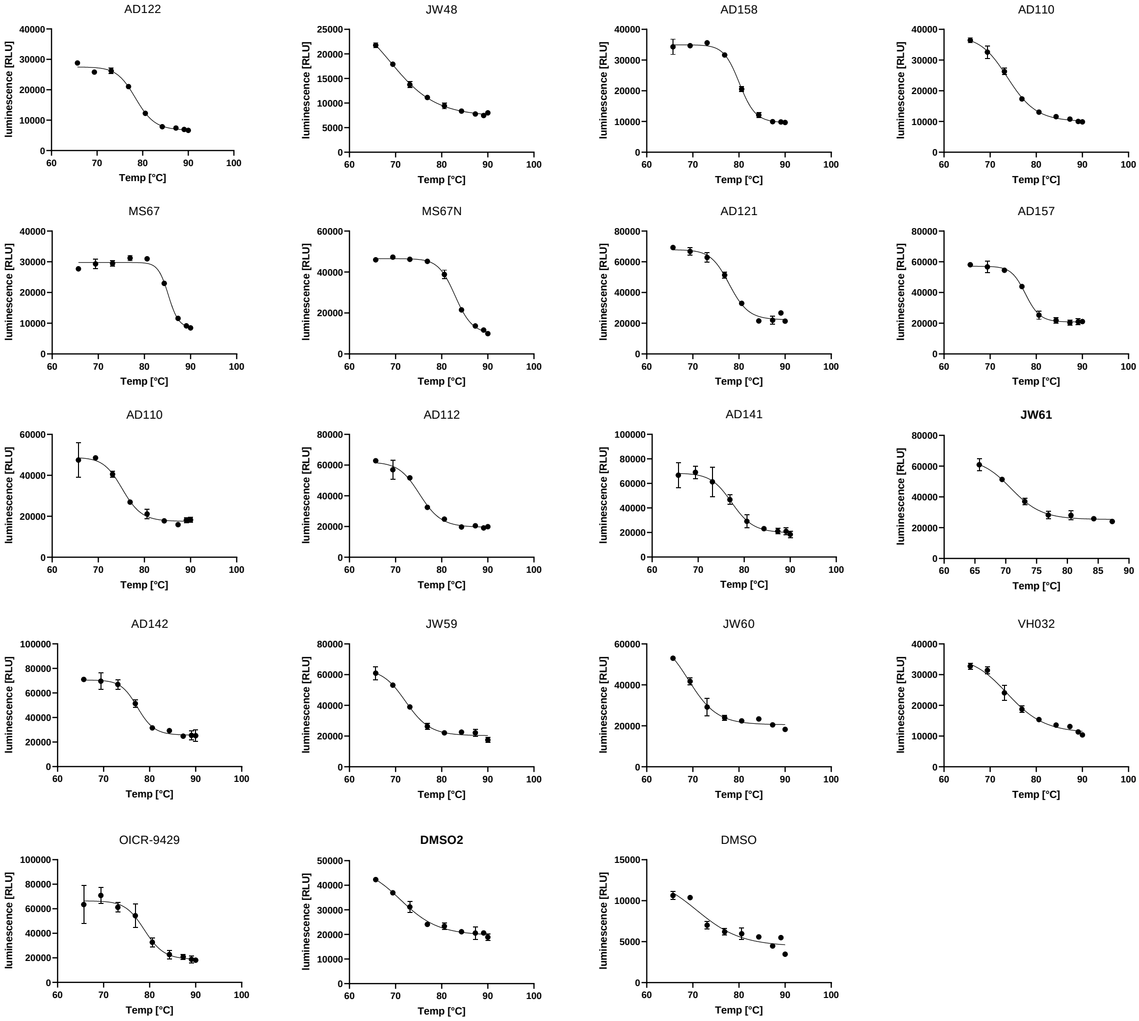


**SI Figure 9: Intact HiBIT-CETSA results plotted as raw data.** Data is expressed as independent biological replicates with error bars depicting the SD. n=2

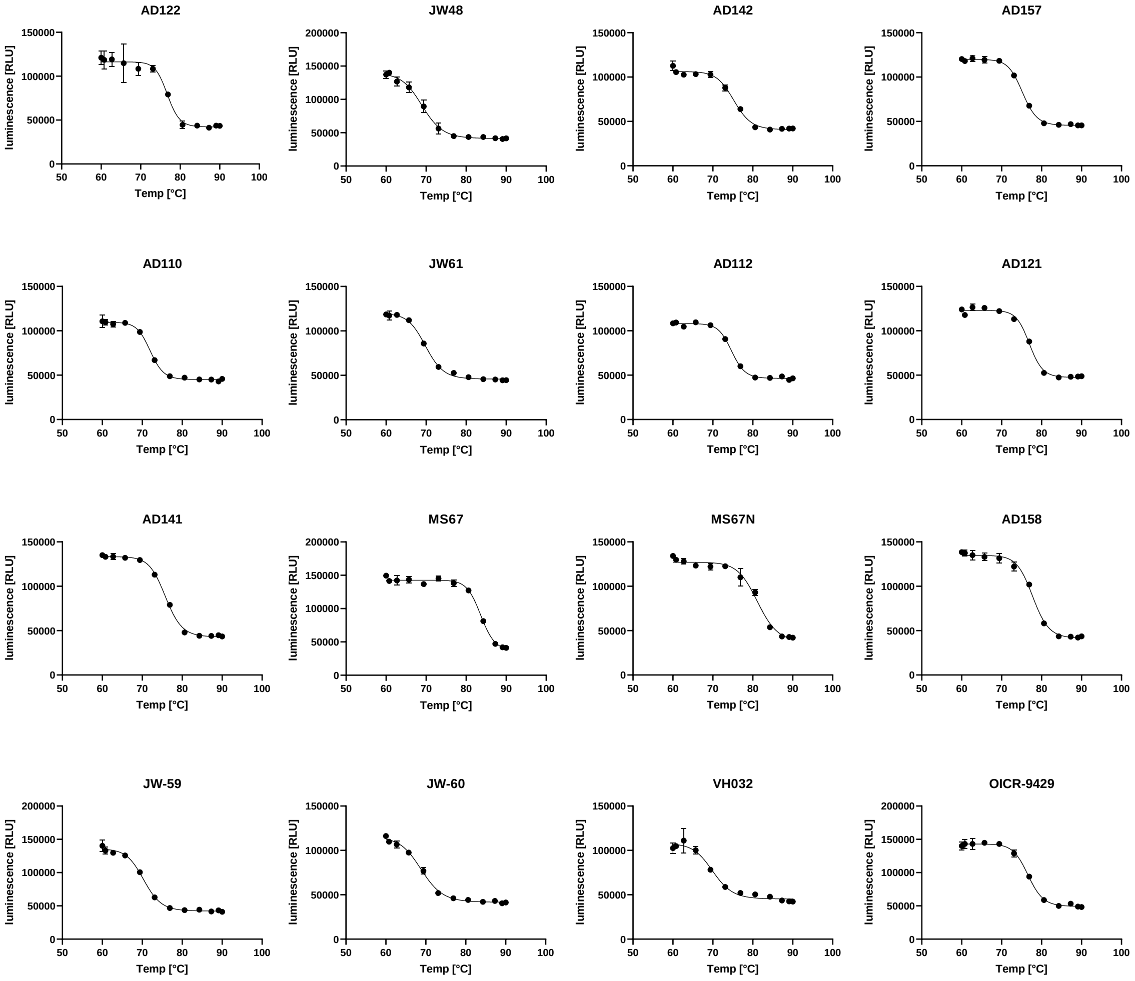


**SI Figure 10: Lysed HiBIT-CETSA results plotted as raw data**. Data is expressed as independent biological replicates with error bars depicting the SD. n=2

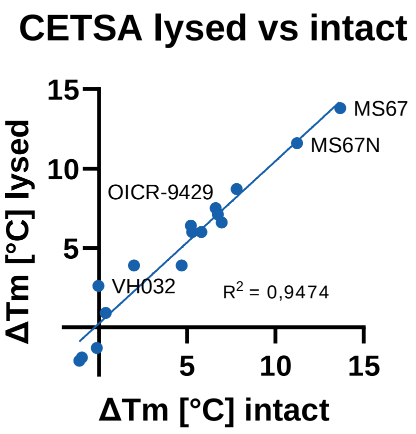


**SI Figure 11: Correlation between lysed and intact cell CETSA results plotted against each other showing correlation with an R2 of 0.95**

**SI Table 2: CETSA results with melting temperatures of intact and lysed cells, respectively**. ΔTm, calculated as melting temperature differences to the DMSO control. ΔTm values published for the *in vitro* WDR5 (Pubmed).

| Compound | CETSA melting temperature | CETSA lysed meting temperature | CETSA ΔTm | Published[1] ΔTm |
|---|---|---|---|---|
| DMSO | 71.4 | 70.0 | 0 | - |
| AD122 (8g) | 78.5 | 76.8 | 7.1 | 13.2 |
| JW48 (17b) | 69.5 | 69.0 | -1.9 | 5.2 |
| AD142 (8j) | 77.4 | 75.3 | 6.0 | 17.0 |
| AD157 (8f) | 77.8 | 75.2 | 6.4 | 10.4 |
| AD110 (8i) | 75.3 | 72.0 | 3.9 | 3.5 |
| JW61 | 70.1 | 69.9 | -1.3 | 3.4 |
| AD112 (8d) | 75.3 | 74.7 | 3.9 | 15.6 |
| AD121 (8h) | 78.0 | 77.0 | 6.6 | 9.7 |
| AD141 (8a) | 77.4 | 75.8 | 6.0 | 15.3 |
| MS67 | 85.2 | 83.7 | 13.8 | - |
| MS67N | 83.0 | 81.2 | 11.6 | - |
| AD158 (8b) | 80.1 | 77.8 | 8.7 | 7.7 |
| JW-59 (17a) | 72.3 | 70.4 | 0.9 | 7.0 |
| JW-60 (17c) | 69.3 | 68.9 | -2.1 | 3.6 |
| VH032 | 74.0 | 70.0 | 2.6 | - |
| OICR-9429 | 78.9 | 76.6 | 7.5 | 13.2 |

**NanoBRET cellular cellular target engagement assay:**

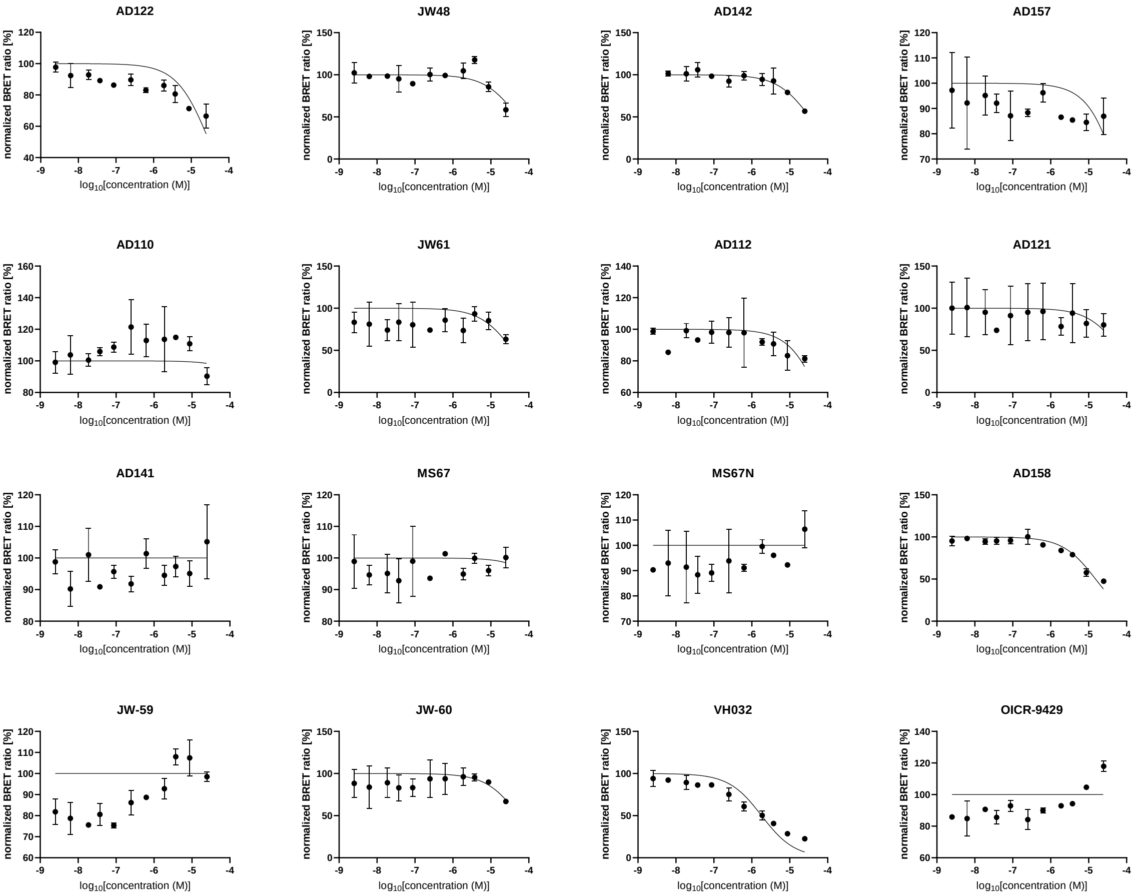

**SI Figure 12: NanoBRET measurements for the compound set against VHL in intact cells**. Data was normalized against a DMSO control. Data is expressed as independent biological replicates with error bars depicting the SD. n=2

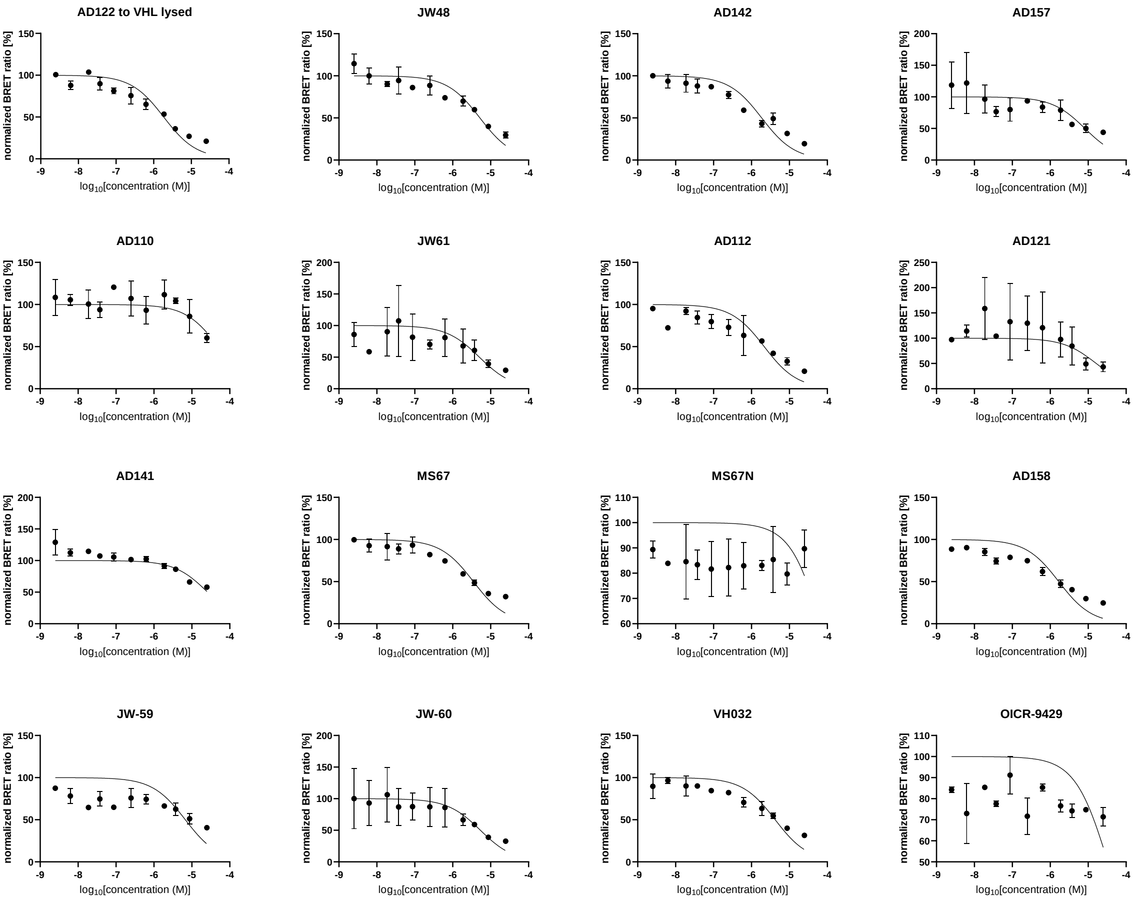


**SI Figure 13: NanoBRET measurements for the compound set against VHL in lysed cells**. Data was normalized against a DMSO control. Data is expressed as independent biological replicates with error bars depicting the SD. n=2

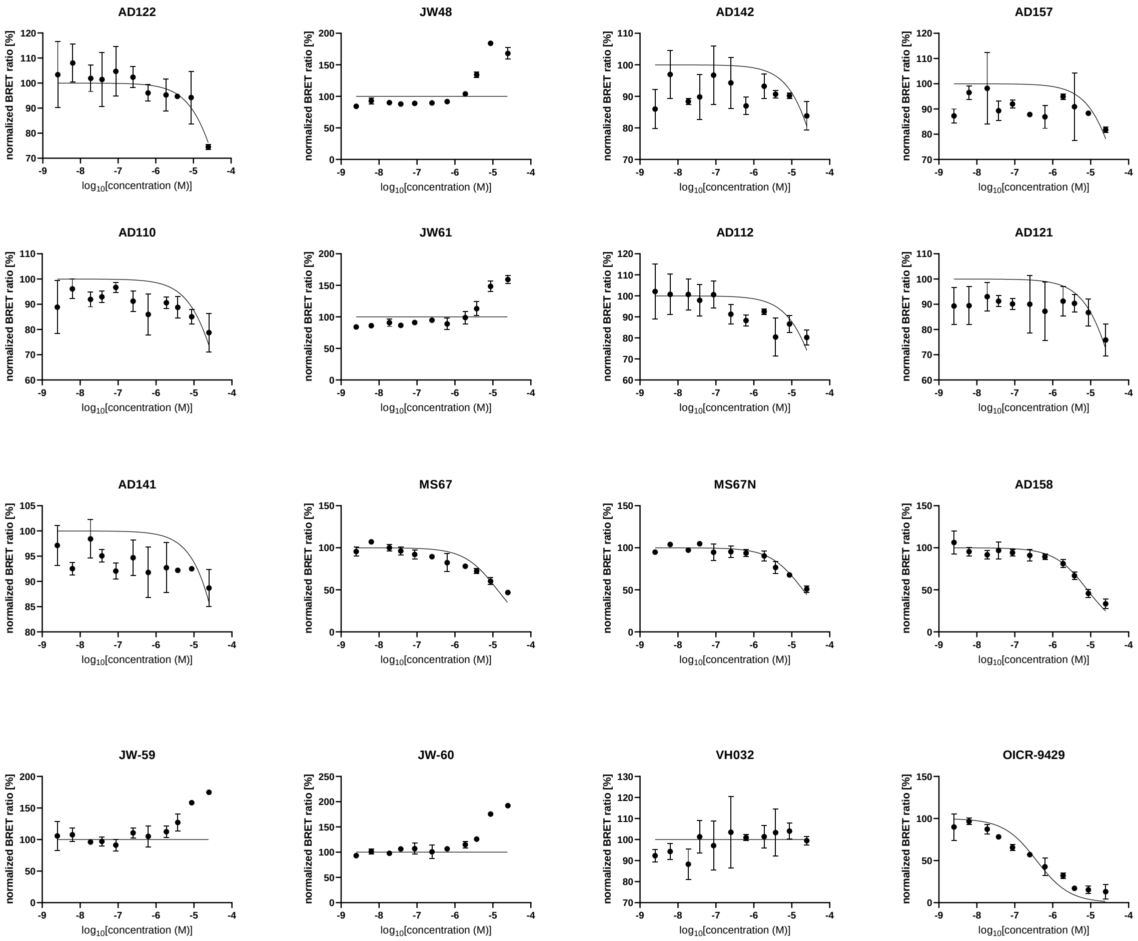


**SI Figure 14: NanoBRET measurements for the compound set against WDR5 in intact cells**. Data was normalized against a DMSO control. Data is expressed as independent biological replicates with error bars depicting the SD. n=2

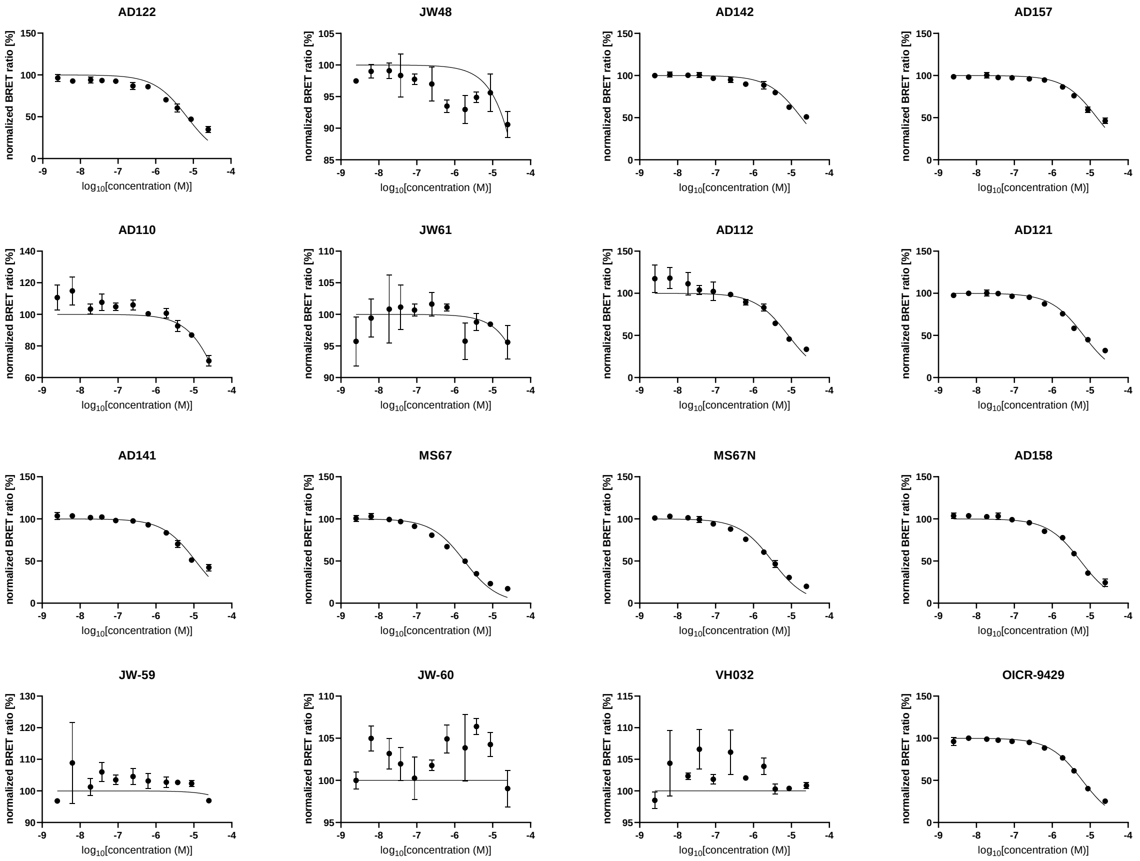


**SI Figure 15: NanoBRET measurements for the compound set against WDR5 in lysed cells**. Data was normalized against a DMSO control. Data is expressed as independent biological replicates with error bars depicting the SD. n=2

**Ternary complex formation:**

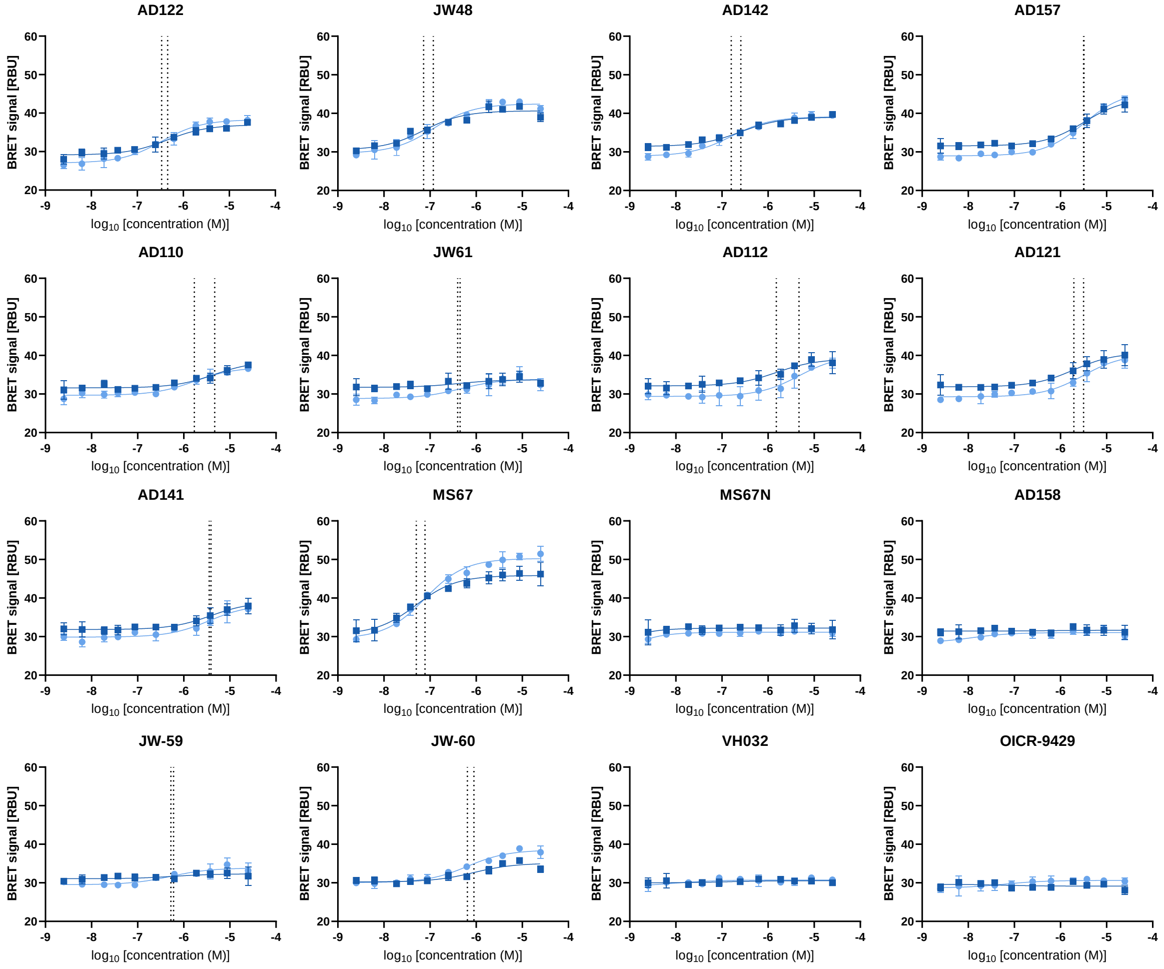


**SI Figure 16: Biological replicate curves of the ternary complex formation assay graphed individually**. Each curve represents one biological duplicate consisting of two technical duplicates with error bars expressing the standard deviation. Dotted lines mark the IC50 value of the individual curves, representing the reproducibility of each ternary complex.

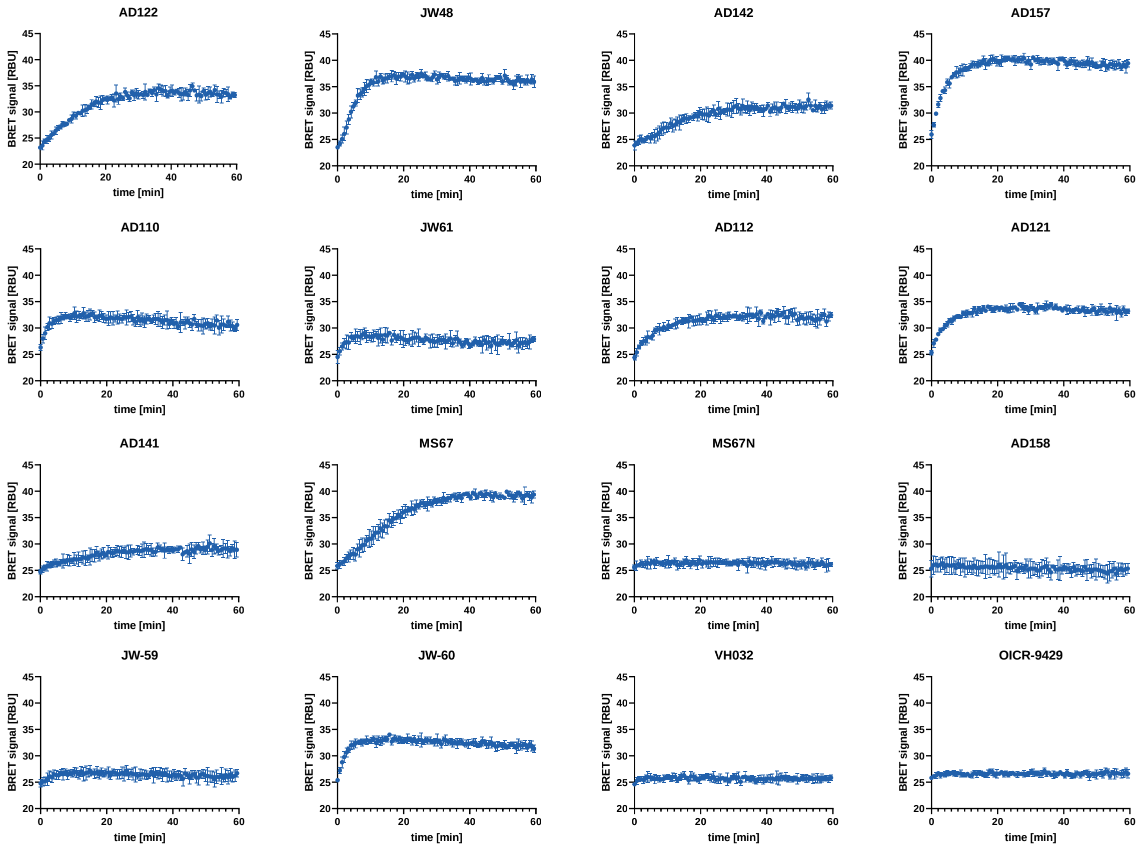


**SI Figure 17: Ternary complex association curves.**

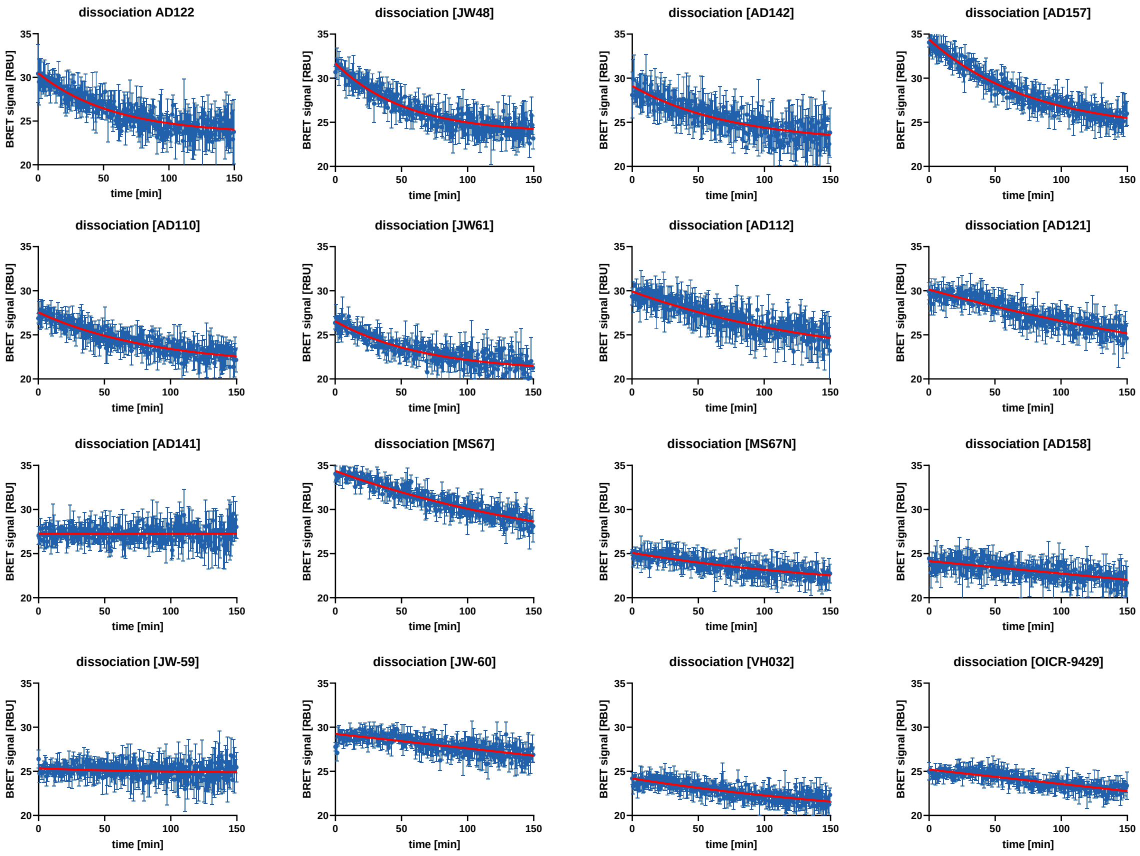


**SI Figure 18: Ternary complex dissociation curves.**

**SI Table 3: Compound concentrations for ternary complex affinity measurements, results and dissociation half times of ternary complexes**

| Compound | Compound concentration used for ternary complex assay [µM] | Ternary complex affinity | Estimated ternary complex association time [min] | Ternary complex half-life [min] | BRET signal intensity [mBU] |
|---|---|---|---|---|---|
| AD122 (8g) | 4.5 | 394.5 ± 56.5 | 36 | 40 | 35 |
| JW48 (17b) | 0.85 | 95.0 ± 22.2 | 17 | 36 | 37 |
| AD142 (8j) | 2.5 | 207.3 ± 49.9 | 50 | 52 | 31 |
| AD157 (8f) | 32.0 | 3234.0 ± 41.0 | 19 | 53 | 40 |
| AD110 (8i) | 46.0 | 3173.0 ± 1457.0 | 10 | 61 | 32 |
| JW61 | 4.0 | 424.1 ± 24.2 | 8 | 47 | 28 |
| AD112 (8d) | 15.0 | 3092.5 ± 1568.0 | 40 | 107 | 33 |
| AD121 (8h) | 20.0 | 2552.0 ± 605.0 | 20 | >150 | 34 |
| AD141 (8a) | 40.0 | 3719.0 ± 189.0 | 50 | >150 | 29 |
| MS67 | 0.75 | 63.8 ± 13.7 | 45 | 134 | 40 |
| MS67N | 0.75 | - | - | >150 | 26 |
| AD158 (8b) | 10.0 | 565.1 ± 43.0 | - | >150 | 25 |
| JW-59 (17a) | 10.0 | 766.3 ± 117.1 | 8 | >150 | 27 |
| JW-60 (17c) | 9.0 | - | 9 | >150 | 33 |
| VH032 | 8.0 | - | - | >150 | 26 |
| OICR-9429 | 8.0 | 394.5 ± 56.5 | - | >150 | 26 |

**Degradation kinetics:**

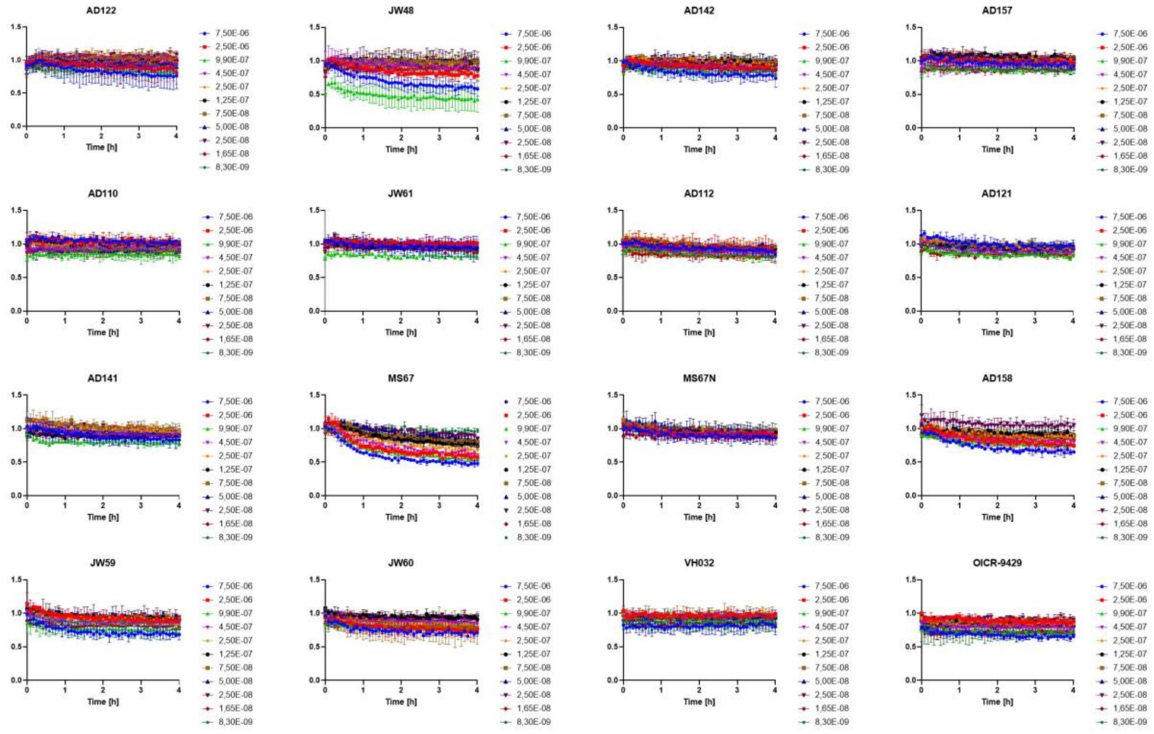


**SI Figure 19: Graphs of the initial 4h of the degradation Kinetics from which the degradation rates were determined.**

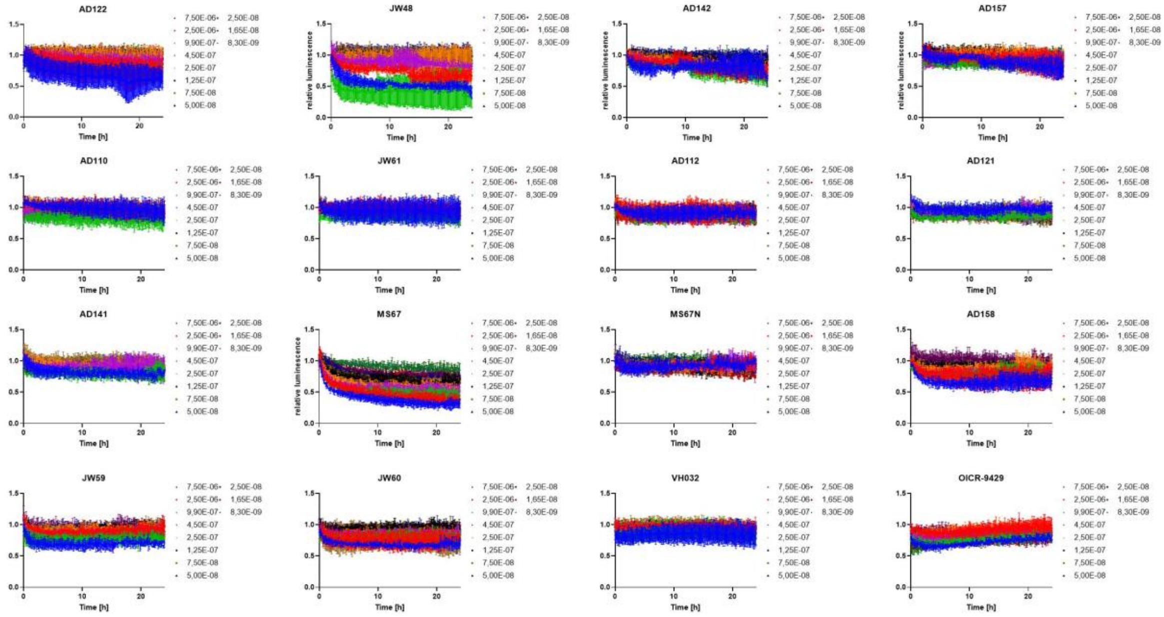


**SI Figure 20: Graphs of the 24h degradation Kinetics.**

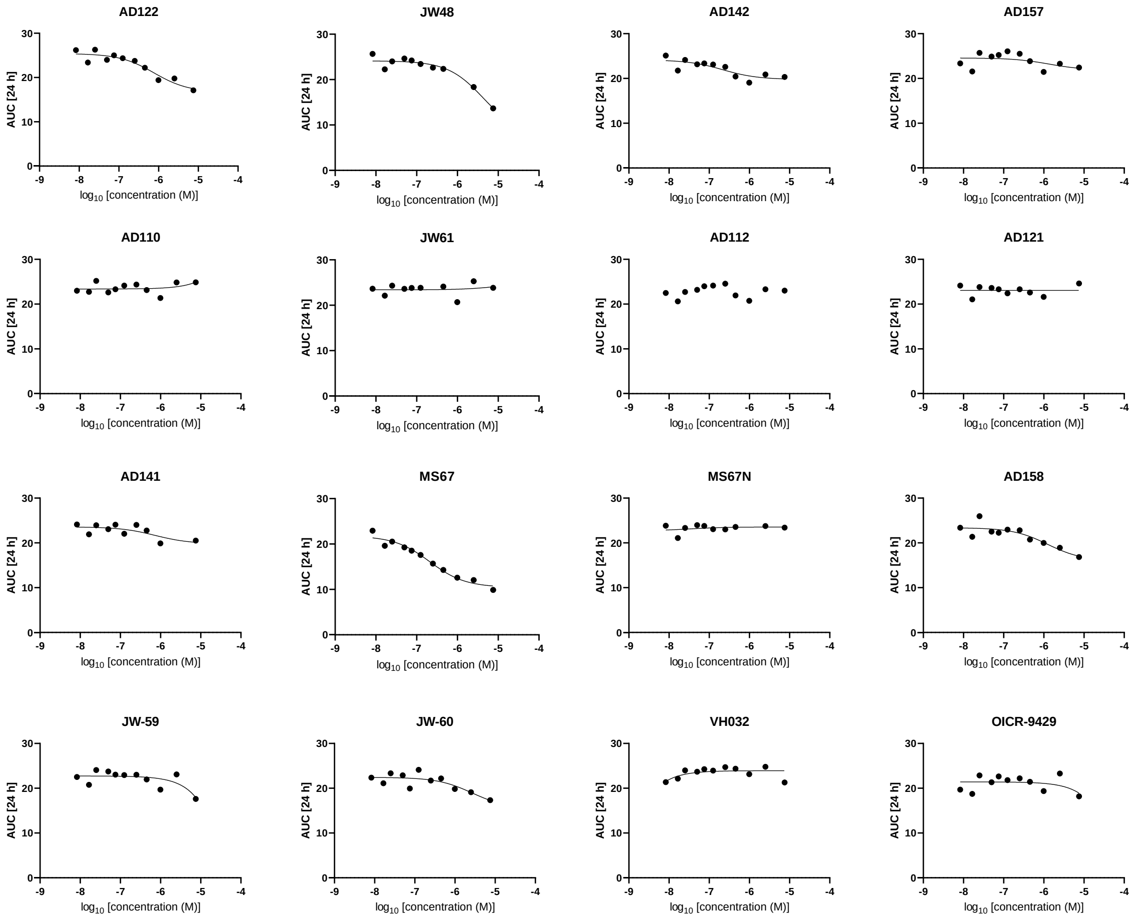


**SI Figure 21: Curves used for DC50 determination over 24h using the AUC.**

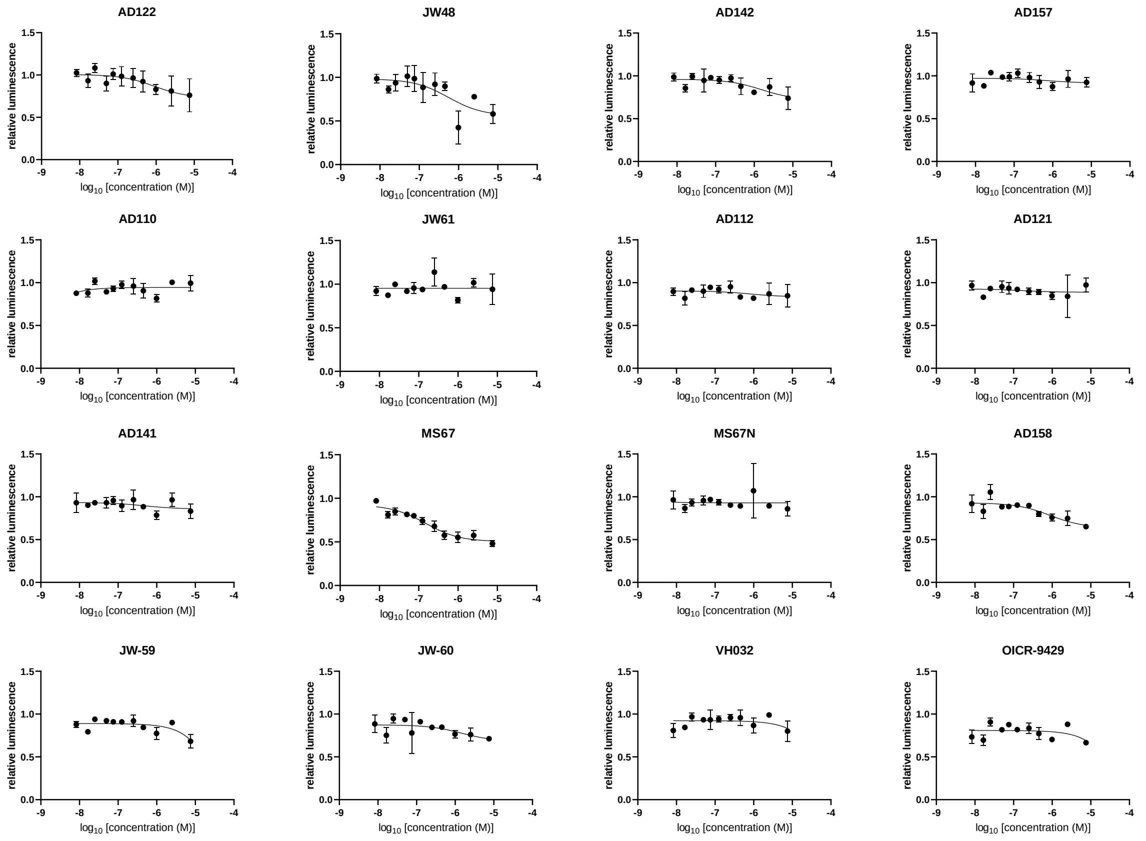


**SI Figure 22: Curves used for DC50 determination at 4h time point.**

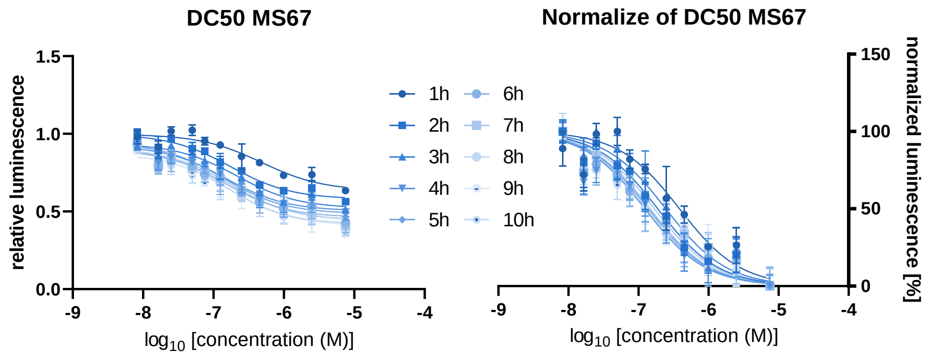


**SI Figure 23: DC50 curves as relative (left panel) and normalised data (right panel) illustrating the time point independency of the DC50.**

**SI Table 4: $DC_{50}$ values for 4h, for the AUC of 24h and the $D_{max}$ between 23 and 24h. $D_{max}$ was calculated as mean from the last 12 measurements from both duplikates with SD calculated.**

| Compound | DC50 (4h) in nM | DC50 (AUC) in nM | Dmax (23-24h) at 7.5 µM in % | Degradation rate (7.5 µM) in $min^{-1}$ |
|---|---|---|---|---|
| AD112 | No | No | 10 ± 6 | 0.48 |
| AD157 | 722 | 1316 | 21 ± 11 | 0.48 |
| AD122 | 896 | 767 | 38 ± 11 | 0.39 |
| AD121 | No | No | 1 ± 4 | 1 |
| AD110 | No | No | 7 ± 11 | 0.11 |

| | | | | |
|---|---|---|---|---|
| AD141 | No | No | 19 ± 4 | 0.65 |
| AD158 | 876 | 1244 | 32 ± 7 | 1.07 |
| AD142 | 1557 | 215 | 28 ± 11 | 1.04 |
| JW-59 | No | No | 30 ± 3 | 1.49 |
| JW48 | 551 | 4275 | 55 ± 6 | 0.75 |
| JW-60 | 1483 | 2675 | 41 ± 5 | 1.3 |
| JW61 | No | No | 6 ± 13 | 1.24 |
| MS67 | 148 | 209 | 69 ± 4 | 1.22 |
| MS67N | No | No | 1 ± 3 | 0.84 |
| VH032 | No | No | No | No |
| OICR-9429 | No | No | No | No |

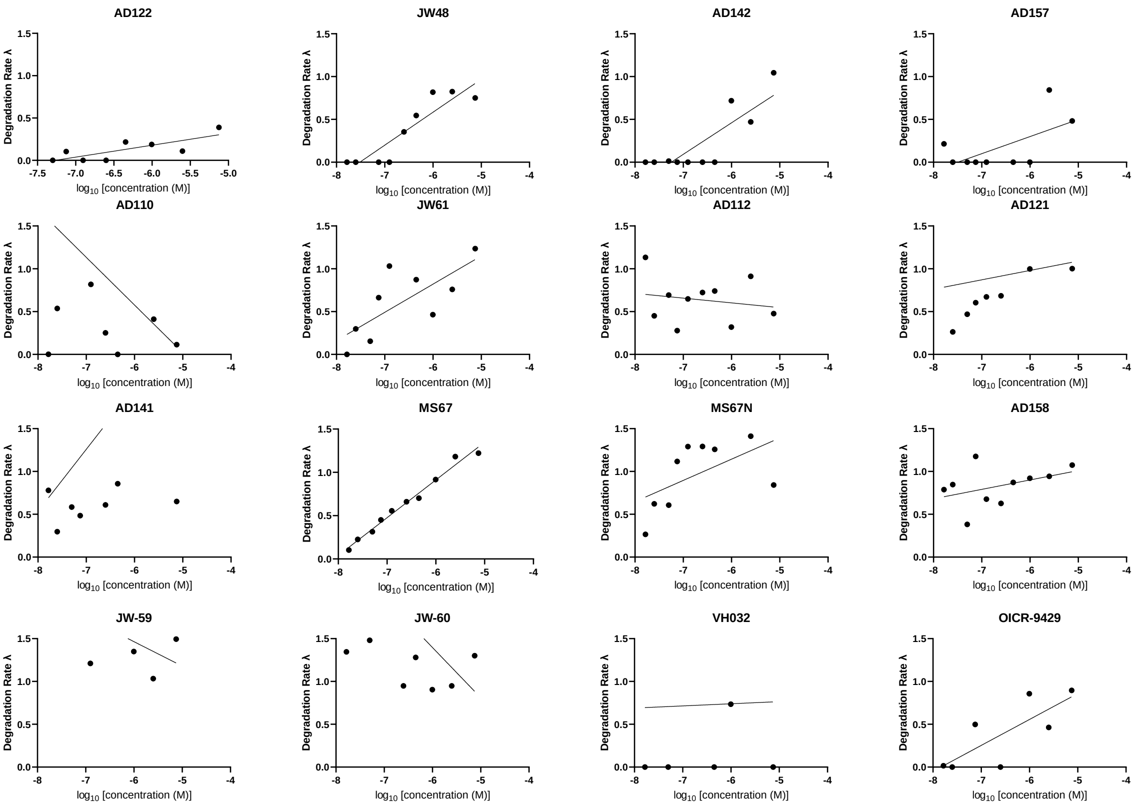


**SI Figure 24: Degradation rates graphed against the compound concentration. Slope and $R^2$ are shown in SI Table 5.**

**SI Table 5: Values for calculated slopes and R2 after correlating the degradation rate against the compound concentration (SI Figure 20). Successful degrader are indicated in red.**

| Compound | Slope | $R^2$ |
|---|---|---|
| AD122 | 0,1408 | 0,618 |
| JW48 | 0,3847 | 0,8308 |
| AD142 | 0,3681 | 0,7002 |
| AD157 | 0,1997 | 0,3762 |
| AD110 | -0,557 | 0,1055 |
| JW61 | 0,3292 | 0,5436 |
| AD112 | -0,05552 | 0,03332 |
| AD121 | 0,1095 | 0,03283 |

| | | |
|---|---|---|
| AD141 | 0,7221 | 0,1072 |
| MS67 | 0,436 | 0,9831 |
| MS67N | 0,248 | 0,3091 |
| AD158 | 0,1099 | 0,1763 |
| JW-59 | -0,2856 | 0,4448 |
| JW-60 | -0,5859 | 0,2045 |
| VH032 | 0,0251 | 0,0002671 |
| OICR-9429 | 0,303 | 0,6072 |

**X-ray crystallography:**

**SI Table 6:**

| **Data collection** | **WDR5-VHL;EloB;EloC-PROTAC** | | | **VHL;EloB;EloC** |
|---|---|---|---|---|
| | **C3-Linker (AD157)** | **C4-Linker (AD122)** | **Aryl-Linker (AD42)** | **JW48** |
| Beamline | X06DA / SLS | X06SA / SLS | X06SA / SLS | X06DA / SLS |
| Wavelength (Å) | 0.99998 | 0.99998 | 0.999998 | 0.999987 |
| Space group | P 1 $2_1$ 1 | P 1 $2_1$ 1 | P $2_1$ $2_1$ $2_1$ | C 1 2 1 |
| Cell dimensions | | | | |
| *a, b, c (Å)* | 46.79, 184.51, 47.95 | 47.70, 190.81, 48.70 | 48.12, 86.69, 196.60 | 73.80, 60.79, 97.23 |
| α, β, γ (°) | 90, 103, 90 | 90, 115, 90 | 90, 90, 90 | 90, 101, 90 |
| Resolution (Å)* | 184.51-2.80 (2.95-2.80) | 47.70-2.50 (2.60-2.50) | 49.15-2.20 (2.27-2.20) | 47.79-2.30 (2.38-2.30) |
| unique observations* | 19494 (2832) | 26976 (3045) | 42660 (3607) | 18781 (1874) |
| *Rpim** | 0.105 (0.459) | 0.062 (0.686) | 0.058 (0.345) | 0.046 (0.477) |
| Completeness (%)* | 100.0 (100.0) | 99.3 (99.6) | 99.8 (99.8) | 99.1 (99.8) |
| Multiplicity* | 2.0 (2.0) | 7.1 (7.3) | 9.1 (9.4) | 7.0 (7.3) |
| mean I/σI* | 6.0 (1.2) | 16.3 (2.1) | 10.1 (2.7) | 9.9 (1.8) |
| CC1/2* | 0.984 (0.707) | 0.990 (0.760) | 0.997 (0.830) | 0.997 (0.752) |
| **Refinement** | | | | |
| *Rwork / Rfree* | 0.203 / 0.279 | 0.243 / 0.293 | 0.199 / 0.247 | 0.223 / 0.261 |
| No. of atoms | 5077 | 5028 | 5522 | 2784 |
| protein | 4948 | 4932 | 5065 | 2704 |
| solvent | 58 | 24 | 334 | 20 |
| heterogen | 71 | 72 | 123 | 60 |
| overall B-factors (Å$^2$) | 48.286 | 67.574 | 34.401 | 57.148 |
| Rms deviations | | | | |
| Bond lengths (Å) | 0.0052 | 0.0062 | 0.0068 | 0.0021 |
| Bond angles (°) | 1.1991 | 1.539 | 1.4193 | 0.7563 |
| Ramachandran outlier (%) | 0.3 | 0.3 | 0.0 | 0.3 |
| **Protein Data Bank entry** | **8BB4** | **7Q2J** | **8BB5** | **8C13** |

*Values for the highest resolution shell are shown in parentheses.

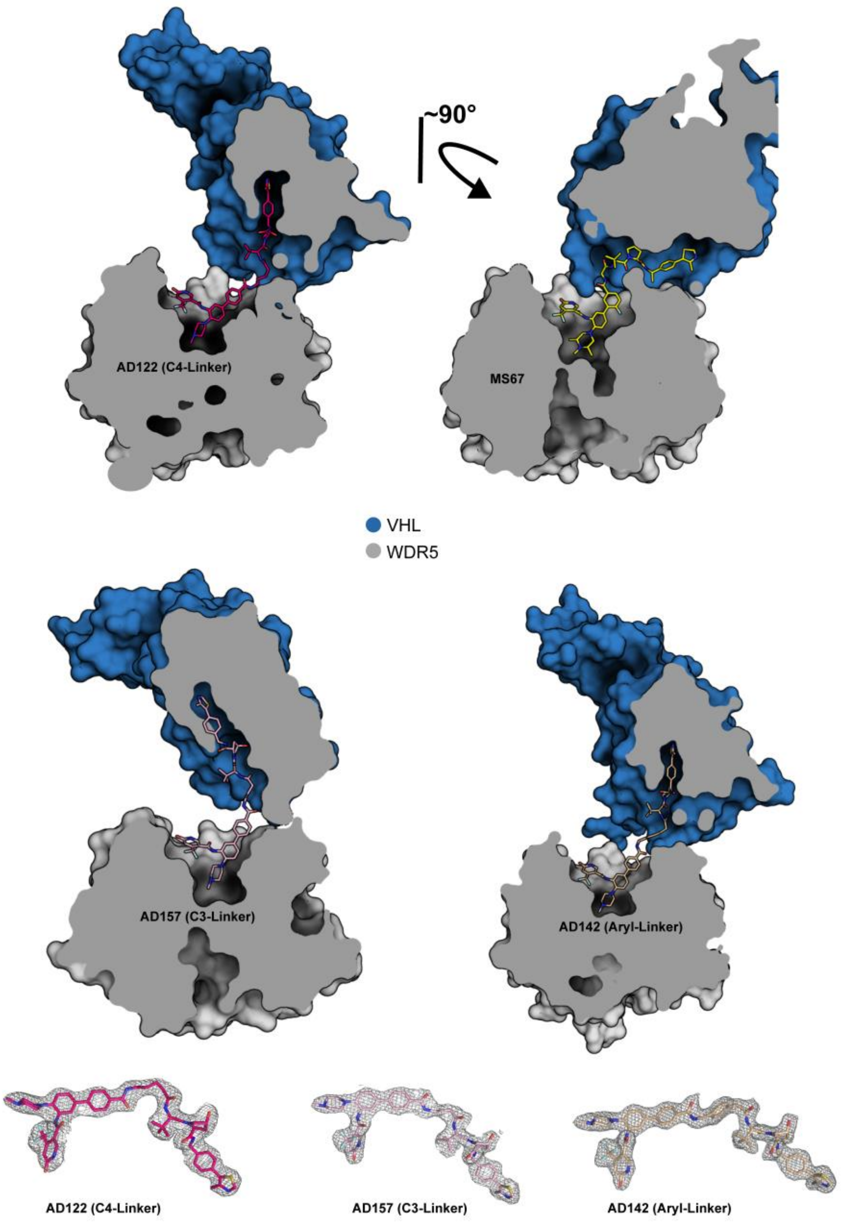


**SI Figure 25: X-ray structures of AD122, MS67, AD157 and AD142. Upper panels show surface representations of the different PROTACs in their binding pockets.** Lower panel displays the electron density map of the bound PROTACs.

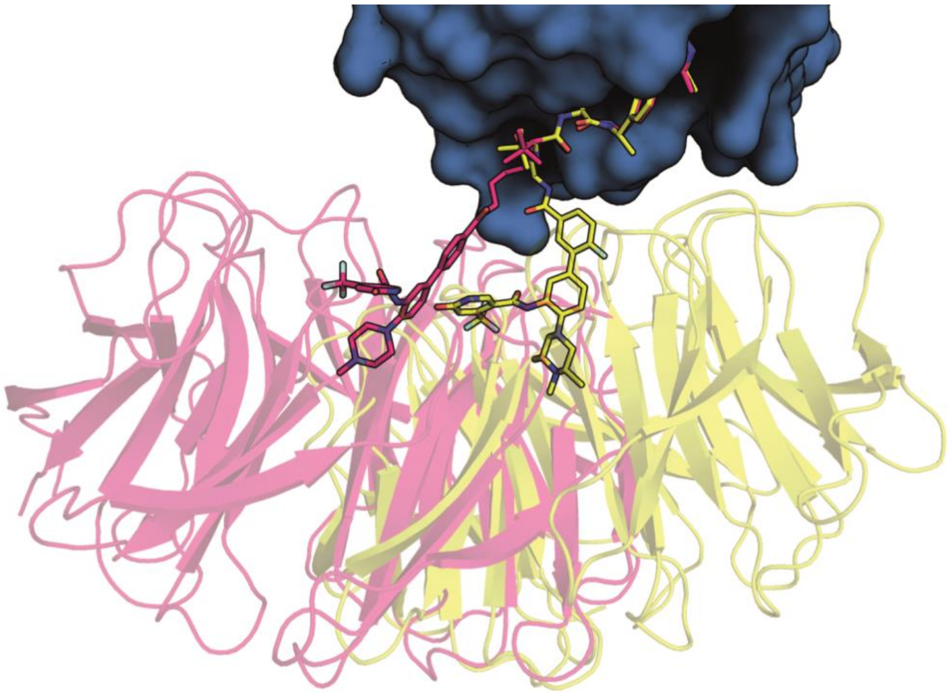

**SI Figure 26: Alignment of AD122 (magenta) and MS67 (yellow) X-ray ternary complex structures indicating the different orientations of WDR5 towards VHL**

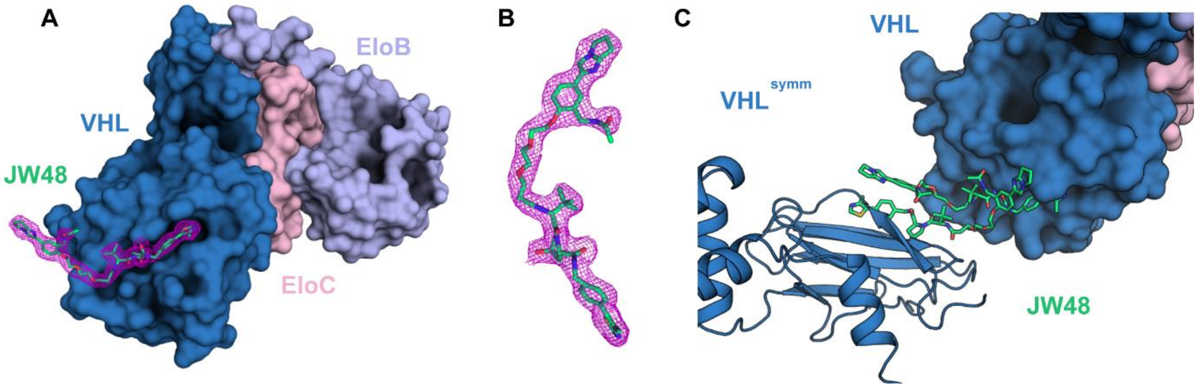


**SI Figure 27: Binary complex crystal structure of VHL-EloB-EloC in complex with JW48 (stick representation).** A) JW48 bound to VHL in surface representation. B) Observed electron density of the partially resolved JW48 countered at 1 σ. C) Symmetry related VHL molecule in the crystal structure (cartoon representation). The orientation of the WDR5 binding part of JW48 is most likely a crystallization artefact.

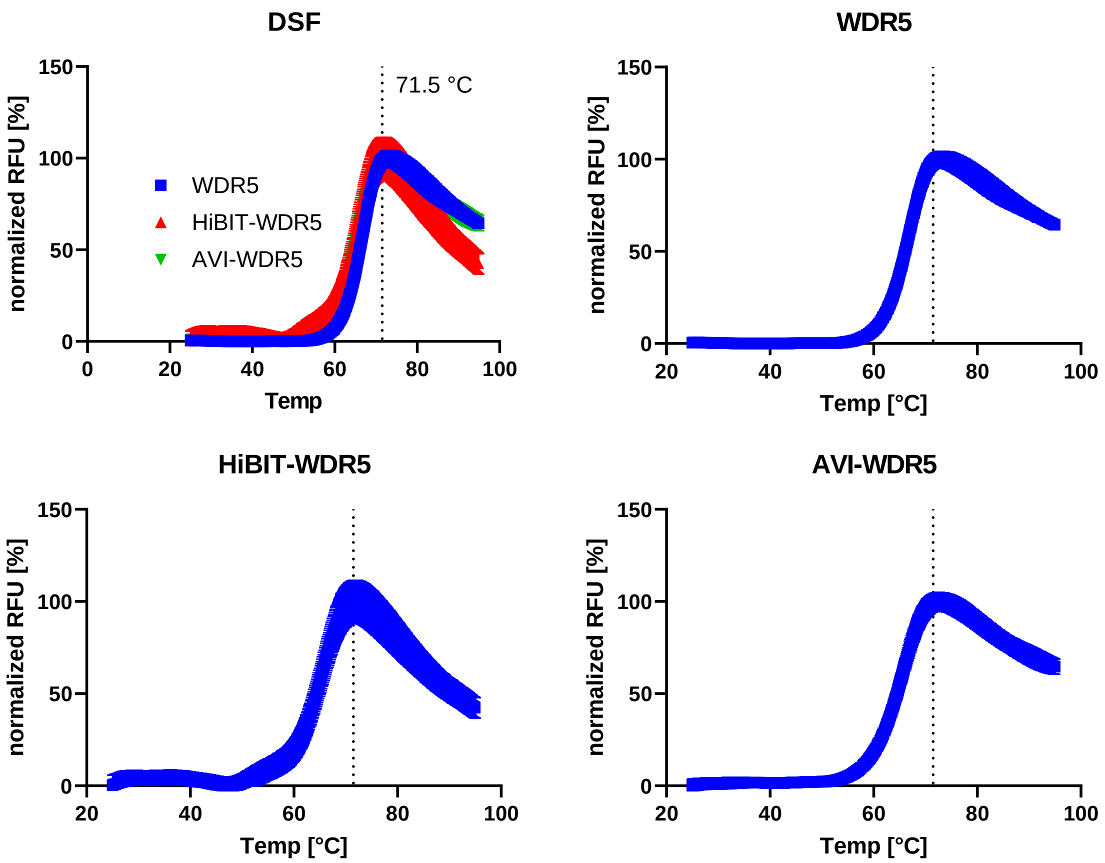


**SI Figure 28: Thermal stability of different purified WDR5 proteins.** Differential scanning fluorimetry of purified WDR5, HiBIT-tagged full length WDR5 or AVI-tagged WDR5 showing an unchanged melting temperature of 71.5 °C indicating unchanged stability upon N-terminal HiBIT tagging. n=4

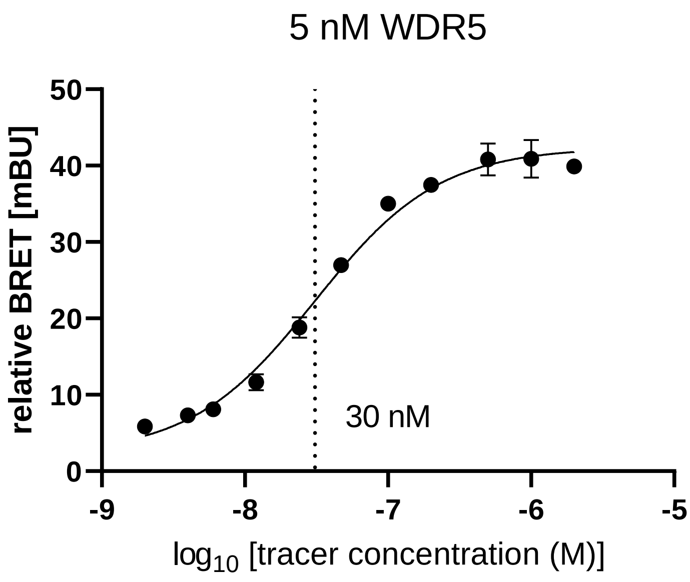


SI Figure 29: Tracertitration curve of the WDR5 tracer towards 5 nM of purified HiBIT-tagged WDR5 protein. The dotted line indicates the tracer $K_D$app of 30 nM. n=2

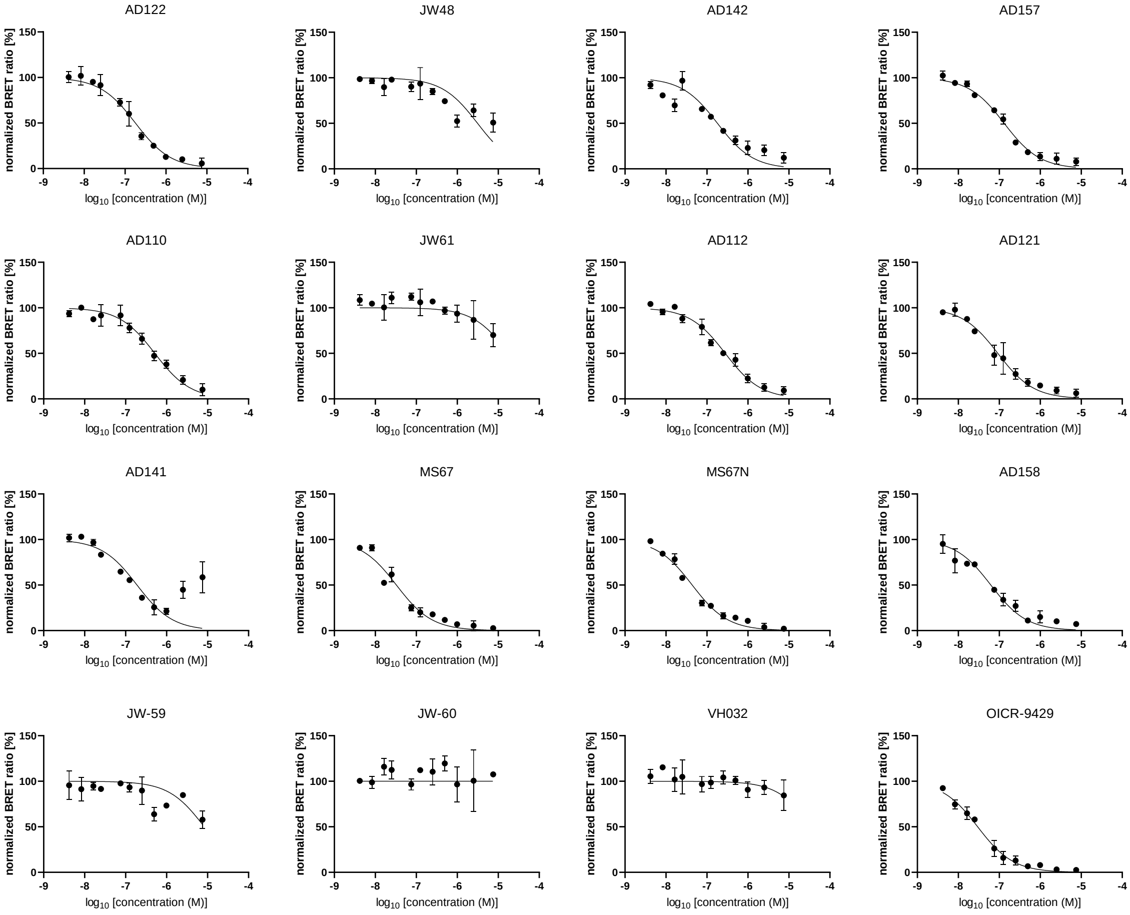


**SI Figure 30: Individual curves of the *in vitro* NanoBRET assay.** HiBIT-tagged WDR5 protein was complemented with the LgBIT and treated with WDR5 tracer and compound. n=2

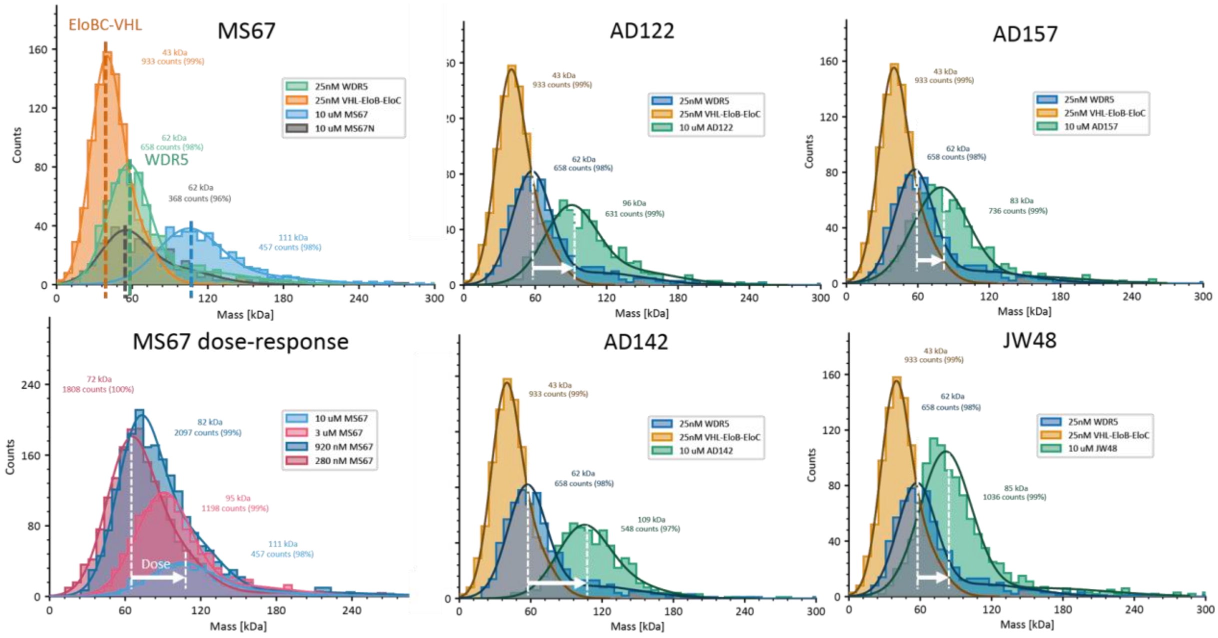


**SI Figure 31: Mass photometry measurements for the individual formed ternary complexes.**

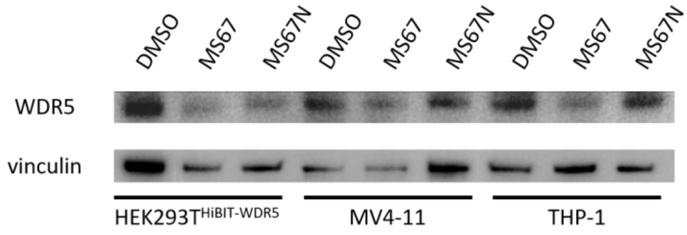


**SI Figure 32: Western blot analysis of the engineered HEK293T cell line in comparison to the wild type AML cancer cell lines MV4-11 and THP-1.** Each cell line was treated with either DMSO or 5 µM MS67/MS67N.
